# The Global Protein-RNA Interaction map of Epithelial Splicing Regulatory Protein 1 defines a post-transcriptional program that is essential for epithelial cell function

**DOI:** 10.1101/2021.05.18.444719

**Authors:** Natoya J. Peart, Jae Yeon Hwang, Mathieu Quesnel- Vallières, Matthew J. Sears, Yuequin Yang, Peter Stoilov, Yoseph Barash, Juw Won Park, Russ P. Carstens

## Abstract

The epithelial splicing regulatory proteins, ESRP1 and ESRP2 are essential for mammalian development through regulation of a global program of alternative splicing of genes involved in maintenance of epithelial cell function. To further inform our understanding of the molecular functions of ESRP1 we performed enhanced crosslinking immunoprecipitation coupled with high throughput sequencing (eCLIP) in epithelial cells of mouse epidermis. The genome-wide binding sites of ESRP1 were integrated with RNA-Seq analysis of alterations in splicing and total gene expression that result from epidermal ablation of *Esrp1* and *Esrp2*. These studies demonstrated that ESRP1 functions in splicing regulation occur primarily through direct binding in a position-dependent manner to either promote exon inclusion or skipping. In addition, we also identified widespread binding of ESRP1 in 3’ and 5’ untranslated regions (UTRs) of genes involved in epithelial cell function suggesting that its post-transcriptional functions extend beyond splicing regulation.

## Introduction

RNA binding proteins (RBPs) have diverse post-transcriptional functions that impact cell properties through regulation of alternative splicing, mRNA localization, mRNA translation, and mRNA stability. While some RBPs are ubiquitously expressed and involved in essential cell functions, many are cell type-specific and thereby fine tune functions that contribute to physiologic roles unique to those cells, such as neurons (Conboy, 2017; Gerstberger et al., 2014; Hakim et al., 2017; Nikonova et al., 2019; Van Nostrand et al., 2020a). We identified ESRP1 and ESRP2 as paralogous epithelial cell type-specific specific splicing regulatory proteins (Warzecha et al., 2009a; Warzecha et al., 2009b). Studies in epithelial cell lines with *ESRP1/2* depletion and in mouse epithelial cells with *Esrp1/2* ablation showed that ESRPs regulate a global alternative splicing program that is enriched for genes involved in epithelial cell functions including cytoskeletal organization, cell polarity, and maintenance of adherens junctions, and tight junctions (Bebee et al., 2015; Dittmar et al., 2012; Lee et al., 2018; Warzecha et al., 2009a). Furthermore, we showed that *Esrp1* knockout (KO) mice exhibited several developmental defects and perinatal lethality, while mice with knockout of both *Esrp1* and *Esrp2* had substantially greater defects. In contrast, *Esrp2* KO mice had no observed abnormalities indicating that while there is some functional redundancy of these paralogs, ESRP1 is most essential. (Bebee et al., 2015; Lee et al., 2018).

Many RBPs have been shown to regulate multiple steps in RNA processing, and emerging evidence has allocated roles outside of splicing to several splicing factors such as *RBFOX1, QK1, MBNL1, PTBP1/2* amongst others (Hafner et al., 2010; Lee et al., 2016; Masuda et al., 2012; Romanelli et al., 2013). In some instances, the multifunctionality of the RBPs is affected by differential localization wherein alternative splicing leads to the production of RBP protein isoforms with predominantly nuclear or cytoplasmic localization. For example, QKI expresses distinct nuclear and cytoplasmic isoforms that regulate mRNA splicing and translation (Fagg et al., 2017; Hafner et al., 2010; Hall et al., 2013; Wu et al., 1999). *RBFOX1*, another well-studied splicing regulator, also has both nuclear and cytoplasmic isoforms. It has been shown that nuclear RBFOX1 is largely responsible for regulating splicing in neurons, heart, and muscle, while the cytoplasmic isoform regulates mRNA stability and translation (Lee et al., 2016). MBNL1, another RBP with predominantly nuclear and cytoplasmic isoforms, also regulates both alternative splicing, mRNA stability, and mRNA localization (Masuda et al., 2012; Wang et al., 2012). While we previously focused on the role of ESRPs in splicing regulation, we also showed that competing 5’ splice sites at the end of *Esrp1* exon 12 lead to the production of ESRP1 isoforms with distinct nuclear or cytoplasmic localization (Yang and Carstens, 2017). Furthermore, we also showed that the *D. Melanogaster* ortholog of ESRP1, *Fusilli*, also expresses distinct splice variants encoding nuclear and cytoplasmic protein isoforms. These observations suggested that ESRP1 also has a phylogenetically conserved function to regulate post-transcriptional steps in the cytoplasm. However, the targets of ESRP1 in the cytoplasm and its functions in this cell compartment have not been determined.

RNA binding proteins, including ESRP1, execute their regulatory functions through sequence specific binding to RNA. We previously identified a high affinity binding motif for ESRP1 *in vitro* (Dittmar et al., 2012). However, the identification of binding motifs for specific RBPs, while providing valuable insight, does not necessarily reflect *in vivo* binding (Van Nostrand et al., 2020a). Crosslinking immunoprecipitation (CLIP) can be used to interrogate the RBP: RNA relationship *in vivo* and when integrated with data from RNA-seq studies on tissues with modulated expression of RBPs we can infer functional roles. For example in the case of MBNL1, separate roles for both nuclear and cytoplasmic were identified using RNA-Seq data from *Mbnl1* knockdown and MBNL1 CLIP which revealed that it not only bound within introns to regulate splicing, but also had significant 3’UTR binding through which it regulated mRNA stability (Masuda et al., 2012). Similarly for RBFOX2, resolution of the functional roles of the cytoplasmic isoform was achieved by interrogation of using crosslinking immunoprecipitations (iCLIP) with transcriptome profiling (Lee et al., 2016). On a global scale, integrative studies combining CLIP and alternative splicing data acquired for a plethora of RBPs from various cell lines has been used to generate RNA maps of how these RBPs bind and regulate alternative splicing (Yee et al., 2018). We previously showed that the binding motif for ESRP1 is enriched in the downstream intron of exons whose splicing is promoted by ESRP1/2, and in the upstream intron and exon body of ESRP-silenced exons (Bebee et al., 2015; Dittmar et al., 2012; Warzecha et al., 2010). However, proof that this position-dependent regulation was due to direct binding of ESRPs to most regulated exons required identification of genome-wide binding sites. In addition, CLIP is needed for the identification of non-splicing related targets of ESRP1. Therefore, to broadly define the post-transcriptional regulatory network of Esrps *in vivo* we combined RNA-sequencing and enhanced crosslinking immunoprecipitation in the epidermis, where the Esrps are highly expressed. Results from the investigations were used to determine genome-wide *in vivo* targets of ESRP1, confirm the preferred motif for ESRP1 binding, and generate a transcriptome-wide map of ESRP1 binding and regulation.

## Results

### Genome-wide profiling of ESRP1 binding sites with enhanced crosslinking immunoprecipitation

We employed enhanced crosslinking immunoprecipitation (eCLIP) of ESRP1 in mouse epidermis to establish that direct binding of ESRP1 is required for position-dependent splicing regulation and to confirm its binding motif *in vivo* (Van Nostrand et al., 2016). To facilitate immunoprecipitation of endogenous ESRP1 from mouse tissues we used CRISPR/Cas9 to introduce two tandem copies of the FLAG tag at the endogenous N-terminus of mouse ESRP1 (*Esrp1*^*FLAG/FLAG*^ mice)(Lee et al., 2020), an approach previously shown to enable extraction of high-quality binding data from cells (Van Nostrand et al., 2017). The epidermis was separated from the back skin of *Esrp1*^*FLAG/FLAG*^ or wild-type ESRP1 (*Esrp1*^WT/WT^) mice to obtain a pure population of epithelial cells, which were then crosslinked and ESRP1 bound RNAs were immunoprecipitated using an anti-FLAG antibody for eCLIP (Figure 1A) (See methods). After determining the optimal RNase concentration to generate libraries of ∼50nt CLIP tags (Figure S1), we sequenced 6 replicate CLIP samples from the *Esrp1*^*FLAG/FLAG*^ mice with corresponding inputs (see Methods for additional details) to an average depth of 41 million reads per CLIP and 37 million reads per input. As a control, we attempted to generate Flag eCLIP libraries from *Esrp1*^WT/WT^ mice, but these samples required high cycle numbers for library amplification and predominantly produced primer dimer artefacts, and thus were not sequenced. This result suggested that the libraries made using *Esrp1*^FLAG/FLAG^ mice were largely specific. Peaks were identified using Piranha(Uren et al., 2012) for each replicate and pooled to identify high confidence binding sites. This analysis was further supplemented by using PureCLIP (Krakau et al., 2017) on three sets of paired replicates from the *Esrp1*^FLAG/FLAG^ samples to identify binding sites at single nucleotide resolution. Using PureCLIP and Piranha, we observed that ESRP1 crosslinks were most enriched in introns and abundant in coding exons, consistent with the functions of ESRP1 to regulate splicing through binding in exons and/or flanking intron sequences (Figure 1B). We also noted many binding sites for ESRP1 in both 5’ and 3’ UTRs, consistent with a post-transcriptional role for the cytoplasmic isoform of ESRP1.

**Figure 1.**
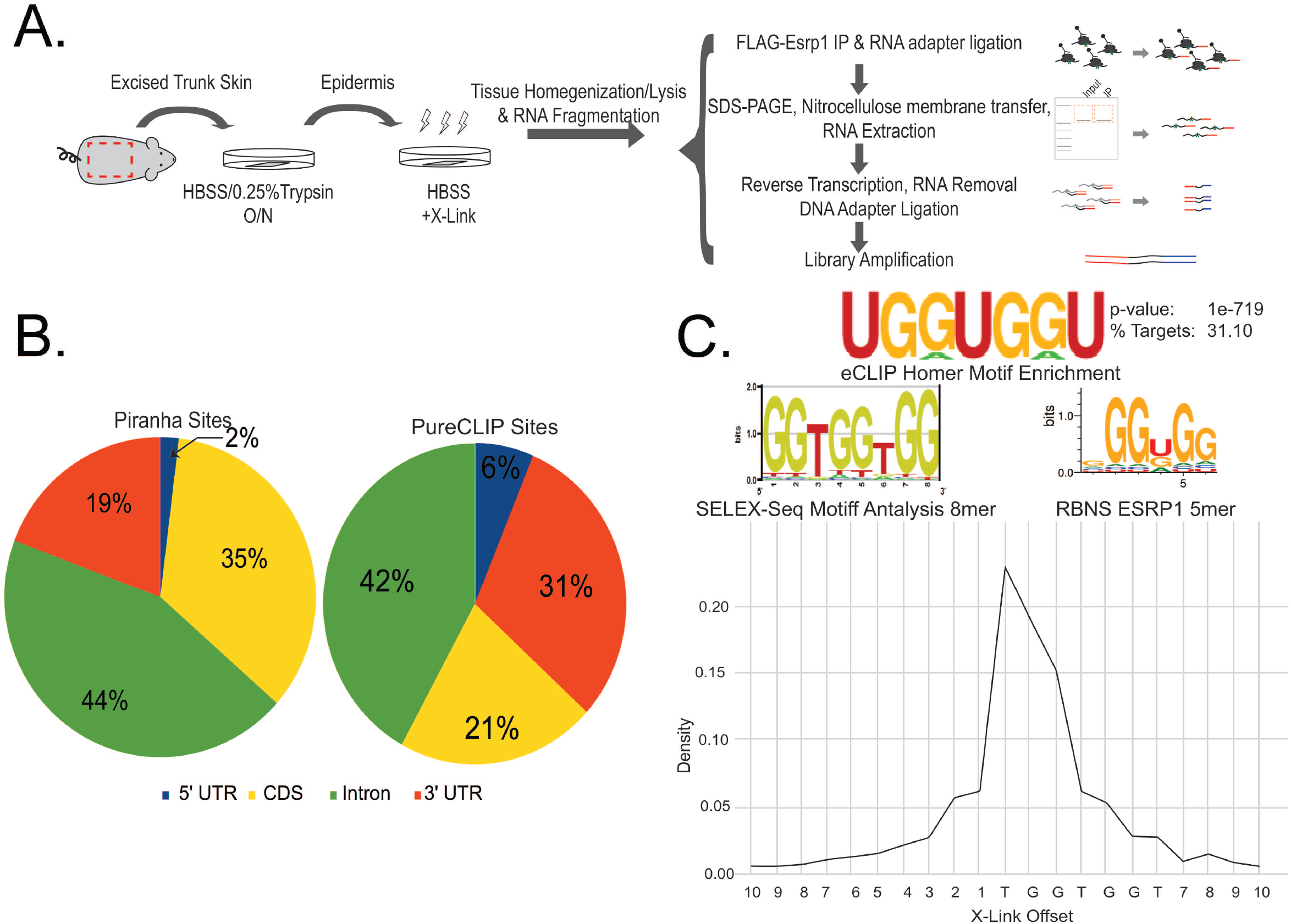
Enhanced crosslinking immunoprecipitation identifies genome wide binding sites for ESRP1. A. Illustration of workflow for enhanced crosslinking immunoprecipitation using Esrp1^FLAG/FLAG^ neonates. B. Pie chart showing of crosslinking sites for ESRP1 identified using PureCLIP and peaks called by Piranha. (Top) WebLogo of de novo motif analysis performed using Homer identifies UGGUGG as most enriched ESRP1 binding motif bound fragments. WebLogo of position weight matrix of 6-mers and 8-mers enriched in ESRP1 bound fragments using Selex-seq (Dittmar et al., 2012) WebLogo of aligned enriched 5mers of ESRP1 (Van Nostrand et al., 2020a). (Bottom) Distribution of crosslinking site frequency relative to the UGGUGG motif shows highest density of crosslinks occur at the first U in the UGGUGG indicating ESRP1 preferentially crosslinks at the second position.

The most enriched motif within the ESRP1 bound RNA fragments consisted of UGG repeats as determined using the HOMER motif discovery algorithm (Heinz et al., 2010). PureCLIP, which facilitates simultaneous peak calling and crosslink site detection at single nucleotide resolution, identified the first position within a UGGUGG motif as the primary site of binding (Figure 1C). This *in vivo* defined binding motif was highly similar to the one we previously identified *in vitro* using SELEX-Seq (Figure 1C) (Dittmar et al., 2012). A highly similar motif was also recently identified using RNA bind-n-Seq in the ENCODE project, further validating the accuracy of ESRP1 binding site determination using eCLIP in *Esrp1*^FLAG/FLAG^ mouse tissues (Van Nostrand et al., 2020a) (Figure 1C).

### Comprehensive identification of transcriptomic changes in the epidermis following inducible ESRP1/2 ablation

To comprehensively identify changes in splicing and total mRNA expression that result from the acute loss of *Esrp1/2* expression we used an inducible knockout strategy to conditionally ablate ESRP1. We previously identified splicing changes in epidermis of mice with germline deletion of *Esrp1* alone or combined deletion of *Esrp1* and *Esrp2* (Bebee et al., 2015). However, these studies used fewer replicates and a lower read depth. It was also possible that some transcriptomic changes in mice with germline *Esrp1/2* ablation were indirect consequences of functional and developmental defects in Esrp ablated epidermis (see below). We therefore used mice with conditional *Esrp1* knockout (*Esrp1*^*flox/flox*^) alleles crossed with transgenic K5rTAand tetO-Cre mice to enable doxycycline inducible ablation of Esrps in neonates by feeding pregnant female mice high concentration doxycycline chow and collecting epidermis from P5 offspring for transcriptomic analysis (Figure 2A). Given some functional redundancy of ESRP2 we used mice with homozygous deletion of *Esrp2*. Littermates were treated as experimental (*K5rTA tetOCre Esrp1*^*flox/flox*^; *Esrp2*^*-/-*^; conditional double knockout (cDKO)) or control (*Esrp1*^*flox/flox*^; *Esrp2*^*-/-*^). We sequenced 4 control and 4 experimental replicates to an average depth of a 150 million reads and assessed changes in alternative splicing and gene expression. Alternatively spliced events identified were congruent with a shift from epithelial to mesenchymal splicing patterns, consistent with a loss of Esrps. We used both rMATs (Shen et al., 2012; Shen et al., 2014) and MAJIQ (Vaquero-Garcia et al., 2016) to assess alternative splicing. Using rMATs we identified 845 splicing changes between control and cDKO epidermis with a |ΔPSI| greater than 10% (FDR≤0.05) (Figure 2B, Table S1). Using a similar stringency of (|ΔPSI|) greater than 10% (P≥0.95) MAJIQ identified a total of 512 local splicing variations (LSVs) that included complex and binary changes in splicing with an absolute change in percentage spliced in (|ΔPSI|) greater than 10% (P≥0.95) (Figure S2A, Table S2). We used semiquantitative RT-PCR to validate over 20 exon skipping events identified by either MAJIQ or rMATS and observe a strong correlation between ΔPSI calculated by using RT-PCR and ΔPSI (r =0.9707, p<0.005; r =0.9704 p<0.005) for MAJIQ and rMATS, respectively (Figure 2C, S2B) with a strong correlation of the reported between rMATs and MAJIQ for shared exon skipping events (r=-0.9872, p<0.005) (Figure S2C). Consistent with a position dependent role of ESRP1 in control of splicing through direct binding, we identified ESRP1 eCLIP peaks in several ESRP1 regulated targets. For example, *Arhgef10L, Arhgef11* and *Lsm14b* exons were previously shown to have large splicing changes in response to ESRP1/2 depletion or ablation and we observed robust peaks for ESRP1 in or near these regulated exons. In the case of *Arhgef11* and *Lsm14b*, we identified ESRP1 binding sites either within the exon or in the upstream intron that were associated with increased splicing upon *Esrp1/2* deletion. In contrast, we identified ESRP1 binding in the intron downstream of an *Arhgef10l* exon that is skipped after loss of ESRP1/2. These observations supported our previous model of an “RNA map” wherein Esrp1 binding upstream of or within an exon induces exon skipping, while binding in the downstream intron promotes exon splicing (Figure 2D) (Dittmar et al., 2012). Using MAJIQ, we were also able to detect additional AS events beyond the five major types: Skipped Exon (SE), Mutually Exclusive Exon (MXE), Alternative 5’ Splice Site (A5SS), Alternative 3’ Splice Site (A3SS) and Retained Intron (RI), such as Alternative Last Exon (ALE) events (Figure S2A). For example we identified a splicing change in *Gpatch2* that altered the ratio of short and long isoforms due to an ALE event (Figure S2D). Overall, the splicing changes we identified with inducible Esrp ablation included many that were previously identified with germline knockout of *Esrp1/2* (DKO) (Bebee et al., 2015), but consisted of a significantly larger dataset of ESRP-regulated events as well as detection of more complex splicing alterations.

**Figure 2.**
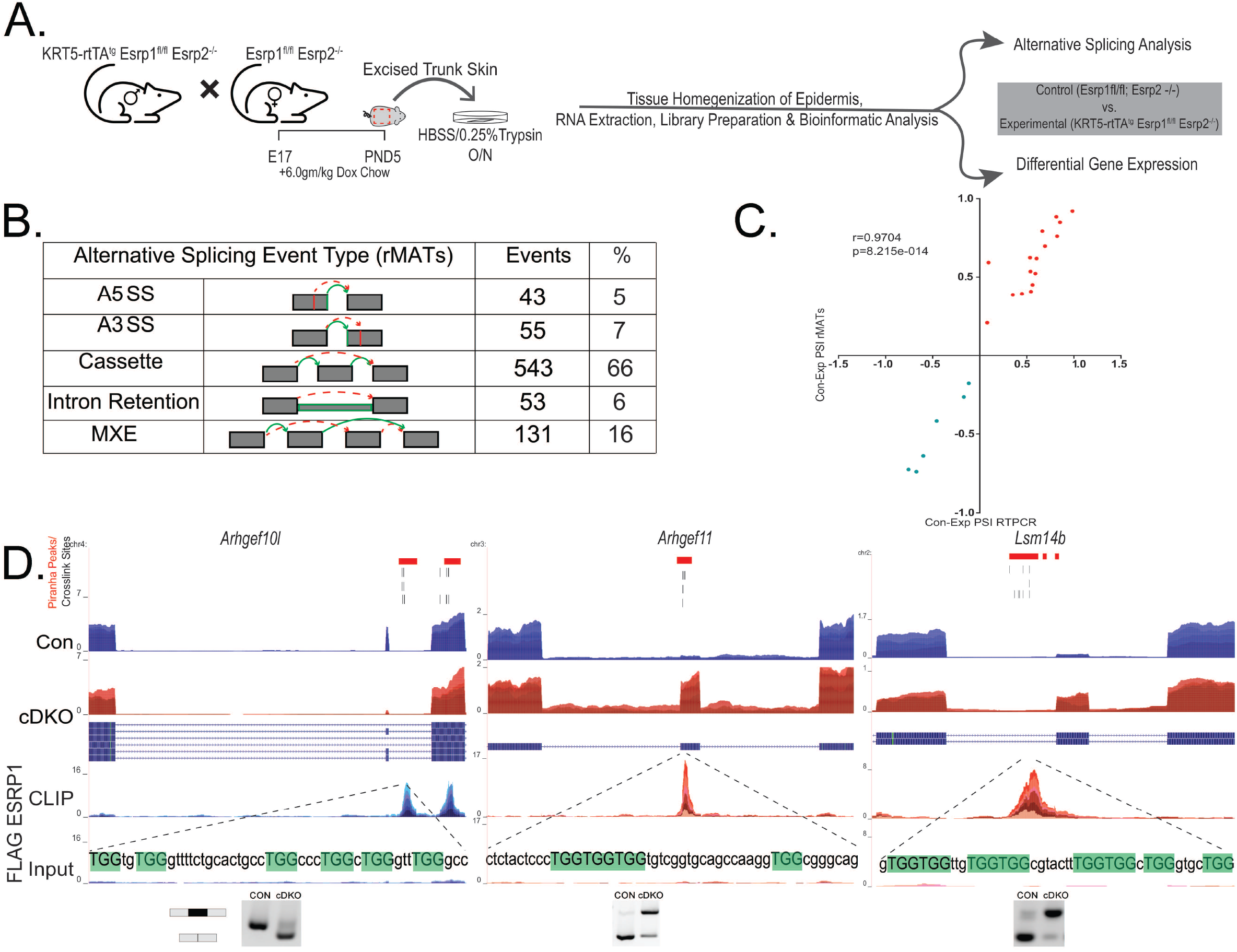
ESRP1 is a direct regulator of alternative splicing. A. Illustration of workflow to generate RNA from control (*Esrp1*^*flox/flox*^; *Esrp2*^*-/-*^) and experimental (*K5rTA; tetOCre; Esrp1*^*flox/flox*^; *Esrp2*^*-/-*^) neonates. B. Number and type of alternatively spliced events identified by rMATs between control and experimental epidermis. C. Correlation plot of ΔPSI estimated by rMATs skipped exon events validated by RT-PCR, with Pearson Correlation r and p-values. Green dots represent splicing events suppressed by ESRP1 and red dots represent splicing events enhanced by ESRP1. D. Genome browser views showing ESRP1 binding peak distribution of pooled replicates for *Arhgef10L, Arhgef11* and *Lsm14b*. Shown are densities in CLIP and size matched Input. Also shown are read densities for transcript levels from RNA sequencing in epidermis ablated of Esrps and control epidermis. UGG motifs are in uppercase and highlighted in green. Positions of crosslink sites from PureCLIP for three replicate pairs and Piranha peaks are shown at top.

### The positions of ESRP1 binding sites globally establish an RNA Map that determines whether it functions to promote exon splicing or skipping

To further characterize the positional effects of ESRP1 binding on splicing at genome-scale we used rMAPS2, a web server which combines differentially regulated AS events data obtained from RNA-seq as determined by rMATs with peaks called from eCLIP-seq data using Piranha (Hwang et al., 2020; Uren et al., 2012). The output from rMAPS2 visualizes the spatial distributions of Esrp1 binding sites near upregulated, downregulated, and background control exons for five major types of AS events: SE, MXE, A5SS, A3SS and RI. This analysis confirmed that ESRP1 binding in the intron downstream of a regulated exon was generally associated with ESRP1 mediated promotion of exon inclusion, whereas ESRP1 binding sites in the upstream intron or within the regulated exon was usually associated with exon skipping (Figure 3A). The CLIP data also demonstrated how position-dependent binding pattern of ESRP1 underlies regulation of more complex changes in alternative splicing. For example, in the case of MYO1B, we noted ESRP1 binding sites upstream of tandem exons (exon 23 and exon 24), that illustrate mechanistically how ESRP1 binding coordinates skipping of both exons (Figure 3B). A similar observation of tandem exon regulation is CD44, where ESRP1 binding sites are present downstream of numerous tandem variable exons case to coordinate splicing of these exons in epithelial cells (Figure S3A). We also noted that three changes in cassette exon splicing in PLEKHA1 corresponded to three splice variant of the same penultimate exon that collectively indicated a greater total splicing change in exon skipping than was calculated using each three variants separately (Figure 3C). To confirm that ESRP1 regulated events are direct targets, we initially examined a total of 543 cassette exons (SE exons) identified by rMATS with |ΔPSI| ≥ 10% and FDR ≤0.05%. For 278 ESRP1-enhanced exons, we identified 47 (17%) that had significant peaks within 250 nucleotides of the downstream intron as determined by Piranha. For 265 exons for which ESRP1 promotes skipping we identified 24 (9%) that had significant peaks in the 250 nucleotides of the upstream exon. We also identified 13 (5%) Esrp1-skipped exons that had peaks within the regulated exon itself (Table S3). While these data fully supported the RNA map of regulation determined by rMAPS2, we noted that the ESRP1-regulated exons in this total list (Table S1, S3) likely consisted of some false positives. In addition we identified multiple examples of definitive CLIP peaks located outside the 250-nucleotide window in adjacent introns. We identified some cases in which the binding sites were more distal, such as an ESRP enhanced exon in ENAH and MYO6 (Figure 4A, S3B). In both ENAH and MYO6, the peak of ESRP1 lies over 1kb downstream of the regulated exon, but the position of ESRP1 binding is consistent with position-dependent regulation by ESRP1. Furthermore, both the regulated exon as well as the region of ESRP1 binding shows high conservation amongst placental mammals and vertebrates. This suggested that a larger percentage of ESRP1-regulated events were regulated by direct binding than indicated by analysis of the peak densities within the 250 nucleotides of the regulated exon. Additional examples of ESRP1 enhanced exons with binding sites in the downstream intron, including far distal sites, in *Slc37a2, Ralgps2, Myo9a, Atp6v1c2*, and *Uap1* are shown in Figure S4. We also illustrate additional examples of ESRP1 silenced exons in *Flnb, Timm17b, Scrib*, and *Magi1* that have ESRP1 binding sites in the upstream intron and/or exon (Figure S5). We examined a more stringent subset of ESRP1-regulated exons that were identified using both rMATS and MAJIQ with an estimated change in |ΔPSI| ≥ 20% (Table 1). For rMATS we further filtered only events with the lowest FDR possible (0) and identified a set of 54 non-redundant simple cassette exons (Table S4). With MAJIQ, we identified a set of 59 manually curated non-redundant cassette exons (Table S4). We extended our analysis to identify any peaks in the entire intron downstream of ESRP1-enhanced exons and the entire upstream intron or exon of ESRP1-silenced exons that were significant peaks by called by Piranha and/or that contained crosslinked site regions identified with PureCLIP. For 39 events that were identified using both rMATS and MAJIQ we identified 29 (74%) with ESRP1 binding sites in a position consistent with the RNA map (Table S4). This analysis suggests that most significant ESRP1-regulated exons are directly regulated. This direct regulation of alternative splicing by ESRP1 was not restricted to skipped exon events but was also observed for mutually exclusive exons such as is the case for FAR1 (Figure 4B) and FGFR2 (Figure S3C). We previously showed that ESRP1 regulates mutually exclusive exons IIIb and IIIc in *Fgfr1, Fgfr2*, and *Fgfr3* (Bebee et al., 2015). While there was complex ESRP1 binding near these exons, a strong binding peak was identified in the intron between exons IIIb and IIIc that is associated with ESRP1-mediated activation of the upstream IIIb exon and repression of the downstream IIIc exon in *Fgfr2*. In contrast, in the case of *Far1*, there is inclusion of either alternate exon 10A or exon 10B (Figure 4B), with binding of ESRP1 appearing to enhance inclusion of exon 10b. In *Fgfr2* and *Far1* both the mutually exclusive exons as well as the ESPR1 binding sites show high conservation. Thus, ESRP1 directly regulates splicing in a position-dependent manner, and this regulation is highly conserved.

**Table 1.**
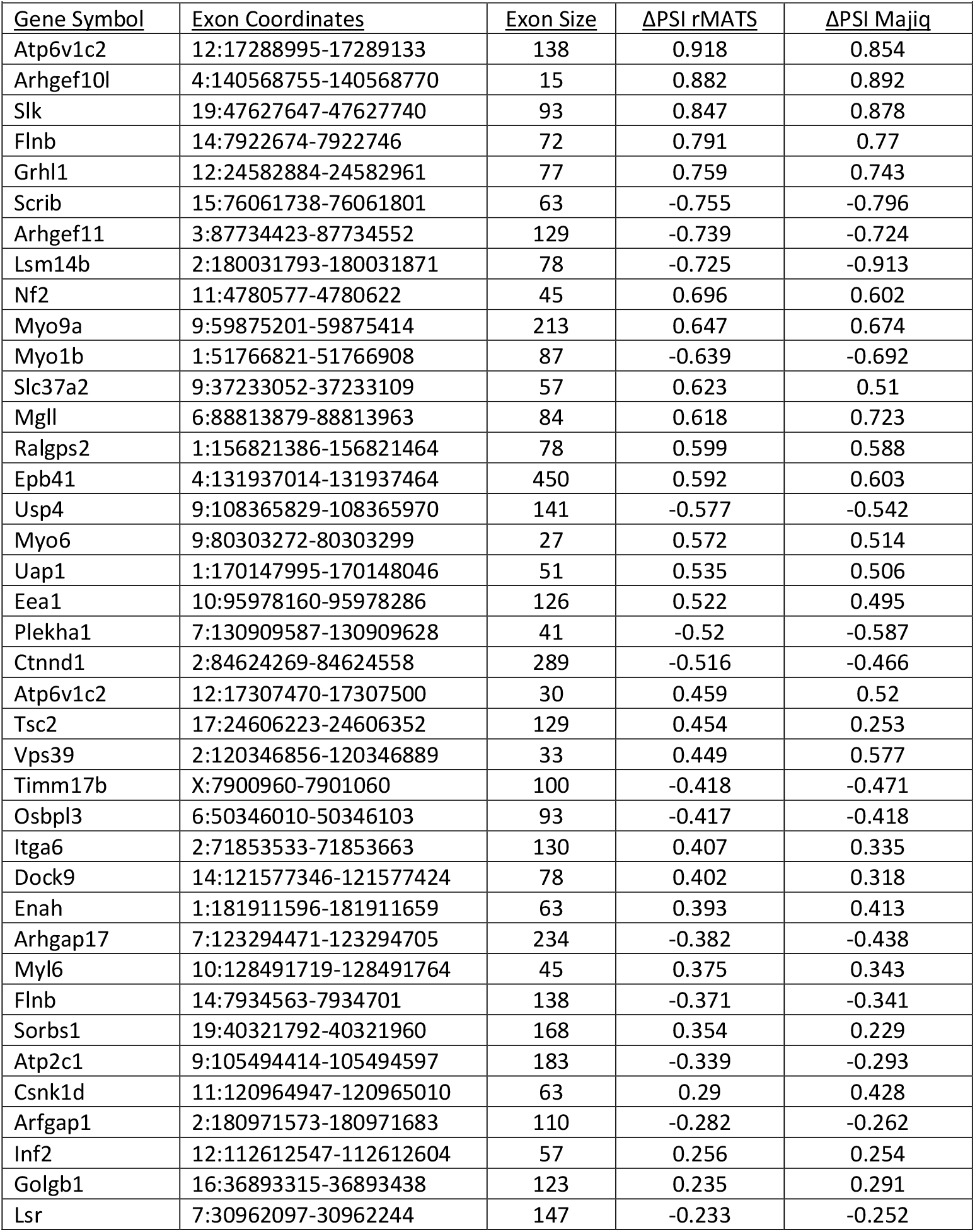
Highly significant changes in cassette exons detected by rMATS and MAJIQ

**Figure 3.**
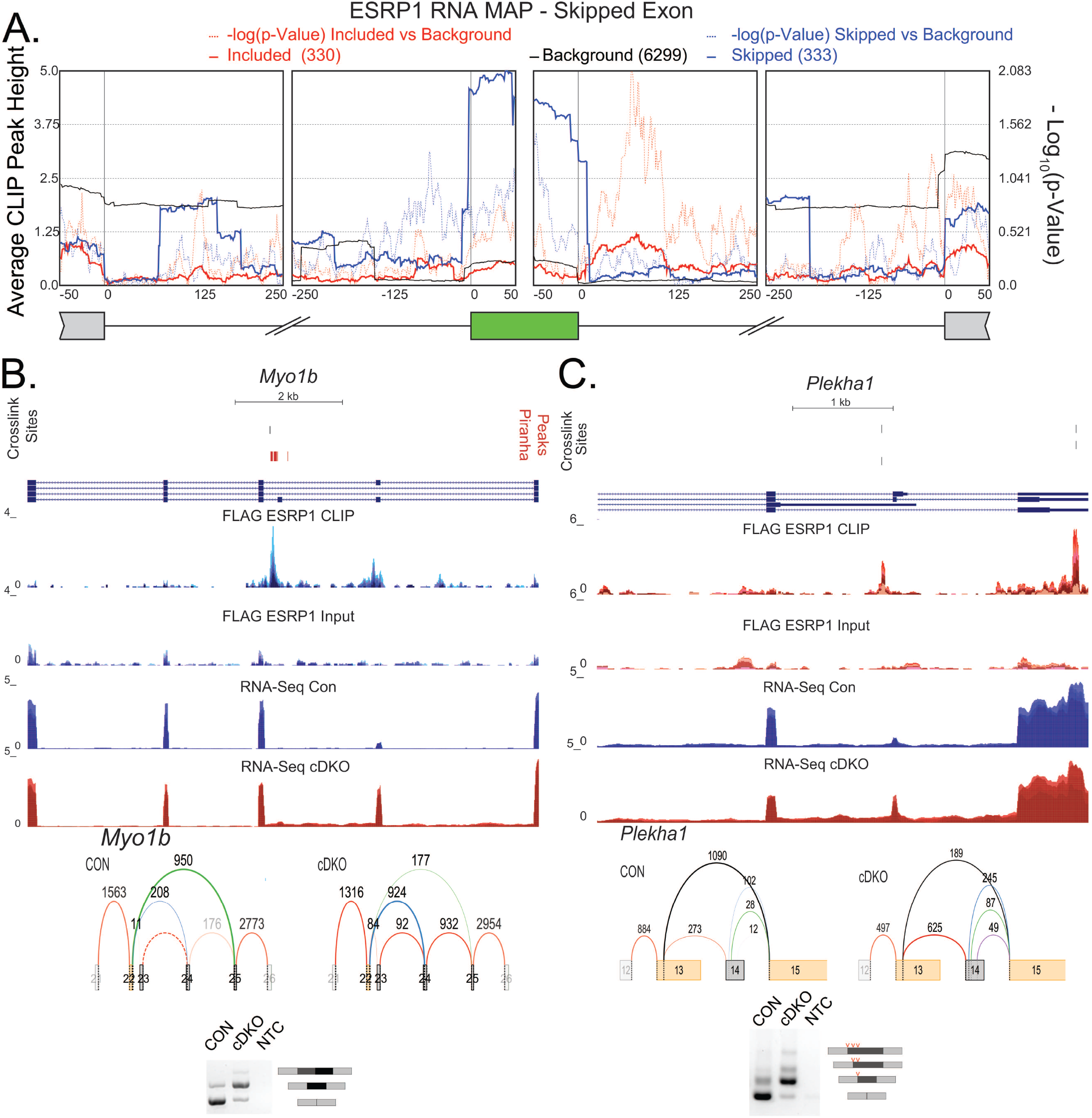
A functional splicing RNA Map for ESRP1. A. Splicing RNA Map of distribution of ESRP1 CLIP peaks on alternatively spliced exons. Solid blue line shows the ESRP1 CLIP densities over ESRP1 downregulated exons, and dotted blue line shows the significance of the peak at each position. Solid red line shows the ESRP1 CLIP densities of ESRP1 upregulated exons, with dotted red line indicating significance (p-value) of the binding at the position shown. B. Voila plots, genome browser views showing ESRP1 binding (CLIP-Seq) and transcript levels (RNA-seq) of for (B) *Myo1b* and (C) *Plekha* and ethidium bromide-stained agarose gel of RT-PCR products of ESRP1 regulated splicing event (red hatches indicate A5SS in *Plekha1)*. Voila plots show differential exon inclusion between Control (CON) and Experimental (cDKO) conditions. CLIP-seq densities show CLIP and size matched input, and transcript levels are from RNA sequence in epidermis ablated of Esrps and control epidermis. Positions of crosslink sites from PureCLIP for three replicate pairs and Piranha peaks are shown at top.

**Figure 4.**
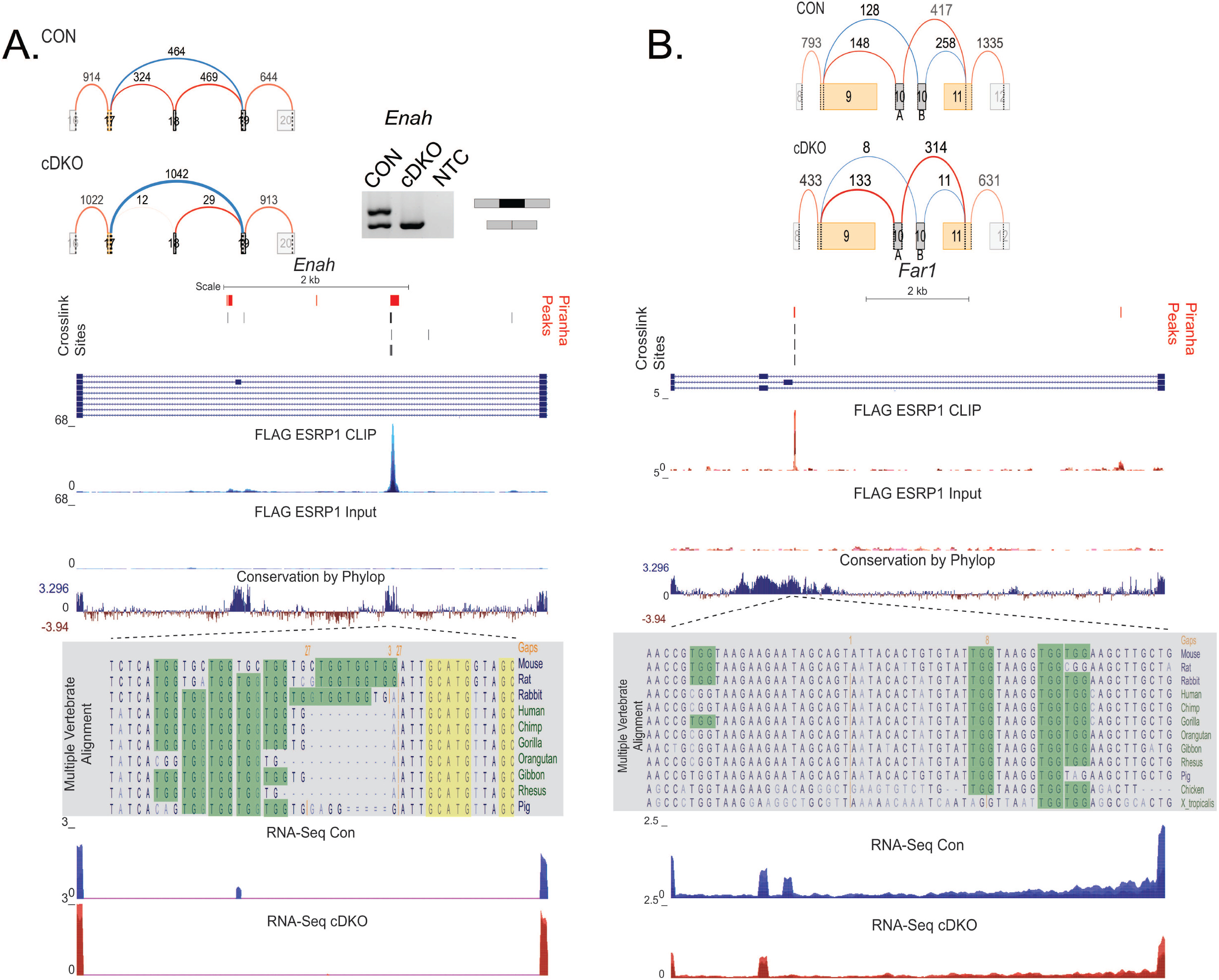
ESRP1 binding and regulation of distal exons. A. ESRP1 binding and regulation of *Enah*. Voila plots and ethidium bromide-stained agarose gel of RT-PCR products of ESRP1 regulated skipped exon splicing event in *Enah*. Genome browser views showing ESRP1 binding (CLIP Seq) and transcript levels (RNA-seq) of for *Enah*. CLIP-seq densities show CLIP and size matched input, and transcript levels are from RNA sequence in epidermis ablated of Esrps and control epidermis. Crosslink sites identified by PureCLIP are shown for three replicate pairs at top of the browser views. Conservation scores showing Placental mammal conservation by PhyloP at the base. Multiple alignment of ∼50 nucleotide regions bound by ESRP1 for several vertebrate species shown. Highlighted in green are UGG motifs recognized by ESRP1, highlighted in yellow is the RBFOX2 motif. B. ESRP1 binding and regulation of *Far1*. Voila plots of ESRP1 regulated mutually exclusive exon (MXE) splicing event in *Far1*. Genome browser views showing ESRP1 binding (CLIP Seq) and transcript levels (RNA-seq) of for *Far1*. CLIP-seq densities show CLIP and size matched input, and transcript levels are from RNA sequence in epidermis ablated of Esrps and control epidermis. Crosslink sites identified by PureCLIP are shown for three replicate pairs and Piranha peaks are shown at top. Conservation scores showing Placental mammal conservation by PhyloP at the base. Multiple alignment of ∼50 nucleotide region under ESRP1 peak for several vertebrate species shown. Highlighted in green are UGG motifs recognized by ESRP1.

### ESRP1 autoregulates ratio of cytoplasmic and nuclear isoforms

ESRP1 peaks were also found within or near alternative 3’ and 5’ splice sites (A3SS or A5SS). We generated a splicing RNA MAP for the A3SS and A5SS for ESRP1 regulated transcripts (Figure 5A, S6). Although we do not observe many examples of A3SS or A5SS events, limiting our ability to conclude strong position-dependent effects, at least one of these has clear functional implications in that we observe a high confidence peak near competing alternative 5’ splice sites of exon 12 of *ESRP1* (Figure 5B, Table S1). We previously showed that use of alternative 5’ splice sites (with identical 5’ splice site consensus sequences) associated with exon 12 of *ESRP1* that are 12 nucleotides apart generates distinct isoforms of ESRP1. Use of the downstream 5’ splice site results in the inclusion the amino acids CKLP that is a component of a nuclear localization signal (NLS) that directs nuclear import of this splice isoform (ESRP1 +CKLP). However, use of the upstream 5’ splice site generates a protein isoform (ESRP1 – CKLP) lacking this essential component of the NLS resulting in cytoplasmic localization (Yang and Carstens, 2017). The observed peak of ESRP1 lies downstream of the A5SS, close to the second competing splice site, and congruent with our RNA Map we would expect that ESRP1 binding in this region would promote use of the upstream 5’ splice site. To assess the functional relevance of the peak near the competing 5’ splice sites of exon 12, we assessed the splicing in both mouse epidermis ablated of *Esrp1* and in human cells lines with knockdown of *ESRP1*. As predicted, we observe that depletion of ESRP1 shifts the cytoplasmic: nuclear ratio towards production of the nuclear isoform (Figure 5B). This finding is consistent with the RNA MAP and indicates an autoregulatory function of ESRP1 in the nucleus to balance the production of both isoforms, in which high levels of ESRP1 leads to inhibition of the downstream A5SS and a shift to the further upstream A5SS leading to production of the cytoplasmic ESRP1 isoform. However, in limiting conditions of ESRP1 the second splice site is not impaired facilitating greater production of the nuclear ESRP1.

**Figure 5.**
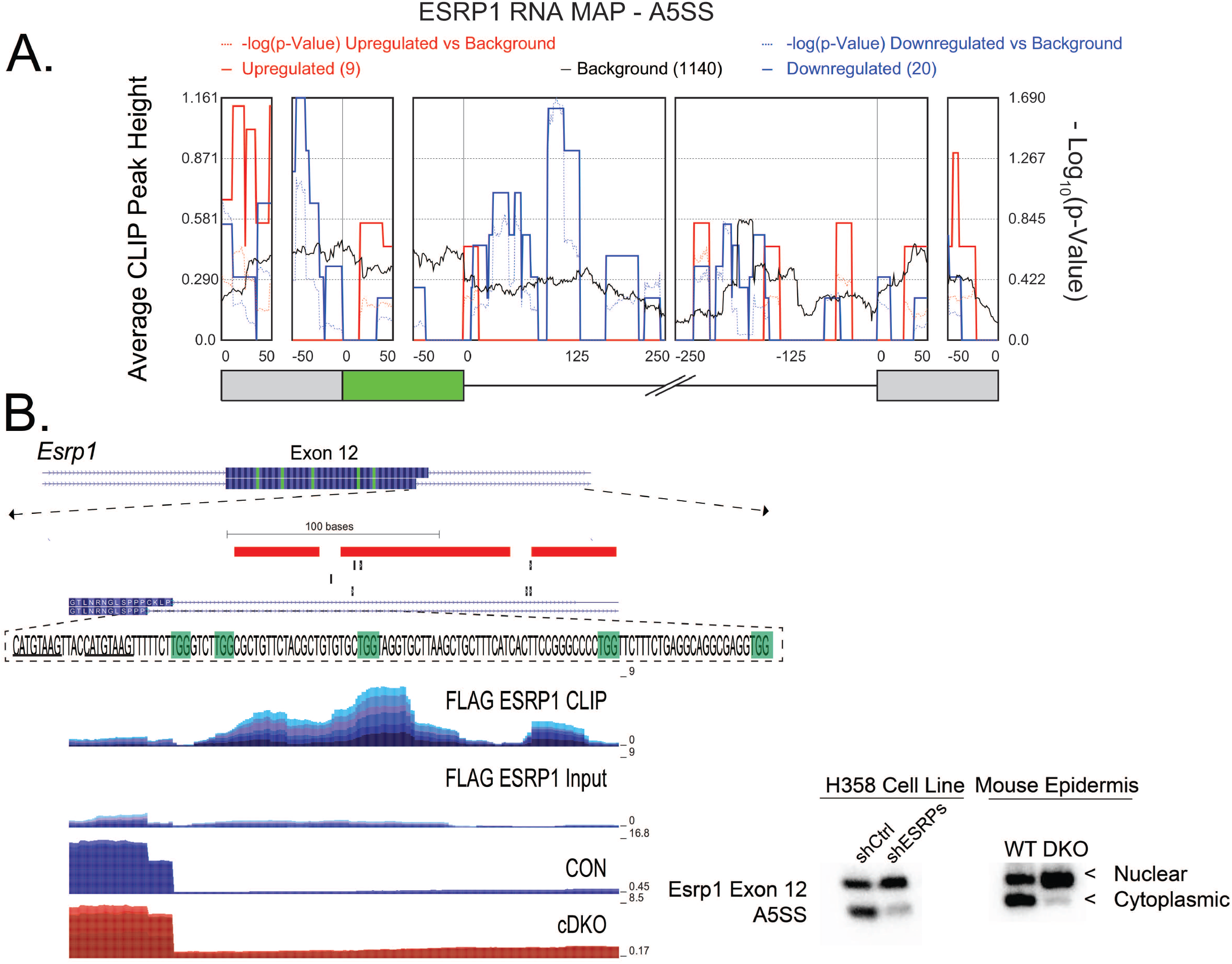
Autoregulation of splicing by ESRP1 balances expression of nuclear and cytoplasmic isoforms. A. Splicing RNA Map of ESRP1 at A5SS. Solid blue line show the ESRP1 CLIP densities at downregulated A5SS, and dotted blue line shows the significance of the peak at each position. Solid red line shows the ESRP1 CLIP densities of ESRP1 upregulated A5SS, with dotted red line indicating significance (p-value) of the binding at the position shown B. (Left) Genome browser view of Esrp1 exon 12 with CLIP seq densities pooled for CLIP and Sm Input showing peak of ESRP1 downstream of 5’ splice site of Esrp1 with crosslink sites and Piranha peaks at top. Highlighted in green text are UGG motifs. Underlined text indicates 5’ splice sites, use of the more distal 5’ splice site results in the inclusion of the CKLP nuclear localization signal. Transcript levels (RNA-seq) show the increase in the use of the more distal 5’ splice site in cDKO conditions compared to control. (Right) Ethidium bromide-stained agarose gel of RT-PCR products of ESRP1 regulated splicing event at exon 12 of *Esrp1* in mouse epidermis ablated of Esrps and control epidermis and H358 cells with or without knockdown of ESRP1.

### ESRP1 binding sites are located in 3’ and 5’ UTRs of genes relevant to epidermal function

We observed substantial numbers of ESRP1 binding sites in 3’ and 5’ UTRs (Figure 1B), suggestive of a function of the cytoplasmic ESRP1 isoform to regulate gene expression in the cytoplasm. We first sought to determine whether ESRP1 binding in UTRs might regulate mRNA stability by evaluating changes in total mRNA levels of these binding targets. However, this analysis was complicated by epidermal barrier defects that result from germline or conditional Esrp ablation (Bebee et al., 2015; Lee et al., 2018). For example, many of the gene expression changes overlap substantially with those that have been described in other mutant mice with epidermal barrier defects such as the knockout models of *Loricrin, Klf4*, and *Grhl3* (Koch et al., 2000; Segre et al., 1999; Yu et al., 2006). These gene expression alterations included components of the epidermal differentiation complex (EDC), including Small proline-rich proteins (SPRR proteins) and Late cornified envelope proteins (LCE proteins) like SPRR1B, SPRR2D, SPRR2E, LCE3C, LCE3E, LCE3F (Bebee et al., 2015). As a consequence, the large changes in total gene expression we observed in *Esrp1/2* KO epidermis may likely be indirect effects of altered barrier function (Kypriotou et al., 2012; Patel et al., 2003). In order to assess a potential direct role for ESRP1 in modulating gene expression changes we used our inducible conditional knockout model of *Esrp1/2* (Figure 2A). We evaluated changes in gene expression using DeSeq2. A total of 1494 genes were differential expressed between neonatal epidermis with Esrps ablated and control epidermis, of which there were 378 genes upregulated more than 2-fold and 456 genes had at least 2-fold or more downregulation and an adjusted p-value <0.05. The gene expression changes observed in the cDKO epidermis was similar to that previously reported in the constitutive ablation of *Esrps* (Figure 6A) (Bebee et al., 2015), which is more consistent with secondary effects of the KO attributable to compensatory changes skin barrier relevant genes as seen in similar mouse models, such as those aforementioned. We explored whether genes with peaks in their 5’ or 3’ UTR were enriched for genes that were altered at the mRNA level. We observed that the vast majority of genes with ESRP1 binding sites in either the 3’ and/or 5’ UTR did not show changes in mRNA levels following inducible Esrp ablation (Figure 5B). Hence, indirect effects of Esrp ablation appear to account for many if not most changes in total mRNA levels.

**Figure 6.**
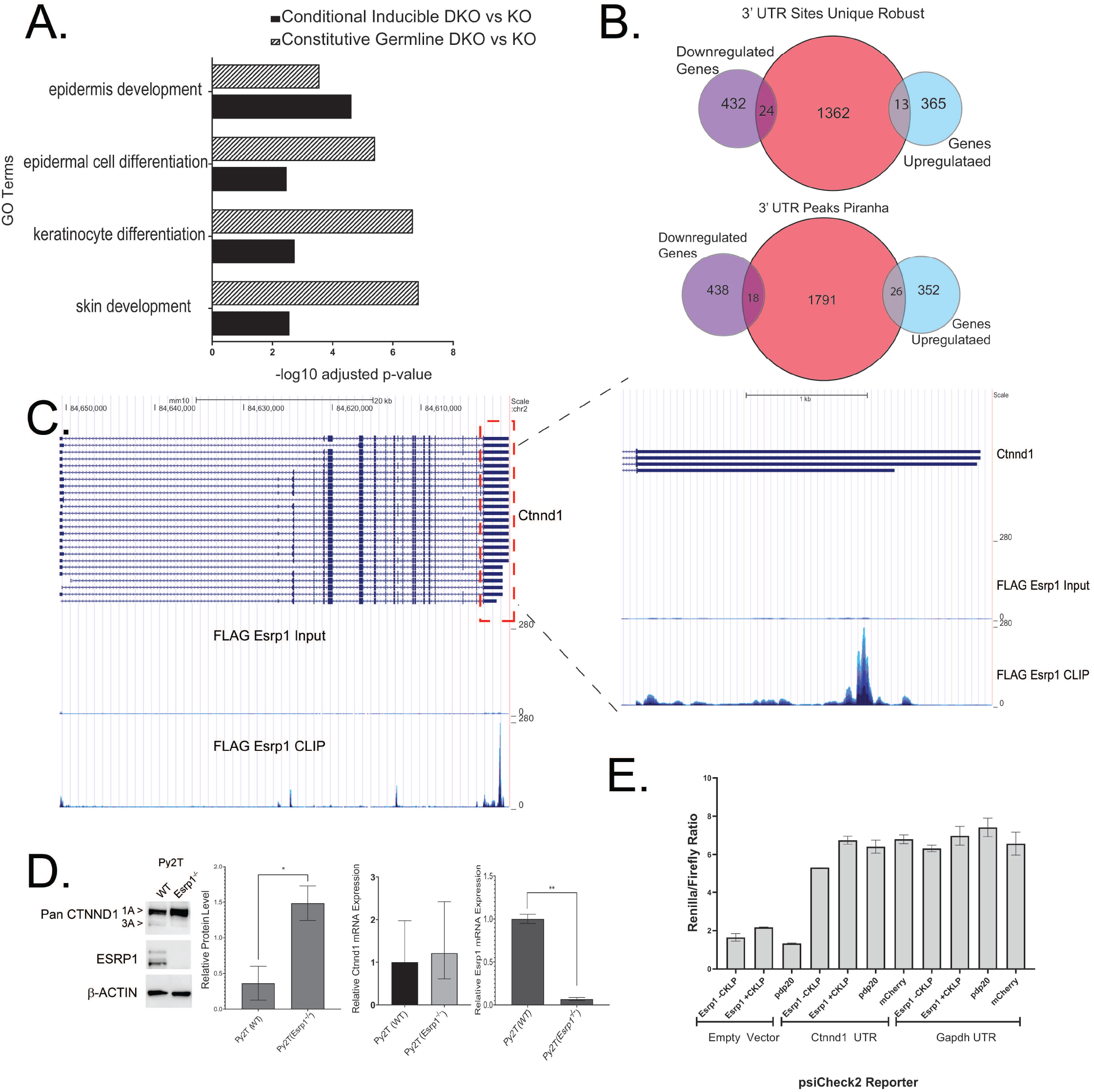
ESRP1 binding in the 3’UTR shows minimal overlap with changes in transcript levels for after *Esrp1* ablation. A. Gene ontology analysis from EnrichR showing the top 4 processes affected in the constitutive Germline knockout of Esrp1 (hatched box) compared to the conditional inducible of knockout *Esrp1* (solid black boxes). B. Venn diagram showing overlap of genes up- or downregulated upon Esrp1 ablation in the mouse epidermis with genes that have peaks of ESRP1 in the 3’UTR as identified by CLIP sites in 2 or more replicate pairs from PureCLIP or from Peaks identified from pooled data from Piranha. C. Genome browser view of Ctnnd1 showing peaks of Esrp1 within the 3’UTR. D. Western blot analysis of Py2T cell lines, *WT* and *Esrp1*^*-/-*^, with *Esrp1* knockout assessing CTNND1. Quantification of the western blots shown, as well as, qRT-PCR analysis of transcript levels for ESRP1 and Pan-CTNND1. E. Luciferase reporter assay comparing effects of Esrp1(+CKLP) or Esrp1 (-CKLP) on *Ctnnd1* UTR and *Gapdh* UTR.

### Esrp1 binds within 3’ UTR of Cadherin binding genes and contribute to altering protein expression

As the majority of genes with ESRP1 binding sites in the 3’ UTR did not show changes in mRNA levels following inducible Esrp ablation (Figure 6B) we were unable to conclusively determine whether ESRPs regulate mRNA stability. To further our assessment, we performed gene ontology (GO) analysis of genes with strong binding peaks in the 3’ UTR using the web server EnrichR (Chen et al., 2013; Kuleshov et al., 2016). We observed that these genes are enriched for Cadherin binding, which is a gene ontology molecular classification for genes that interact selectively and non-covalently with membrane protein, cadherin (Table S5). To assess whether ablation of ESRPs affected protein expression of ESRP1 bound transcripts without changing mRNA expression, we assessed protein levels via western blot of a subset of epithelial targets. We focus on CTNND1 which has crosslinking peaks for ESRP1 in the 3’UTR, Intron, and CDS (Figure 6C). To assess functional consequences of ESRP1 binding in the 3’UTR of CTNND1 we used a cellular model, the Py2T mouse epithelial cell line. We previously used CRISPR/Cas9 technology to generate Py2T *Esrp1*^*-/-*^ clonal cell lines (Lee et al., 2018). We performed western blots in wildtype Py2T cells, and Esrp1 deficient Py2T cells, and observed that as expected there was a reduction in the overall amount of the predominantly epithelial isoform of CTNND1, p120-3. We have previously shown that ESRP1 regulates the expression of the epithelial CTNND1, p120-3 (Warzecha et al., 2009a). Moreover, consistent with direct regulation by ESRP1, we observe ESRP1 binding within the introns upstream of the exons skipped in the p120-3 (Figure S7). However, we also observed an overall increase in total CTNND1 protein expression in Py2T *Esrp1*^*-/-*^cells compared to WT Py2T cells (Figure 6D). We confirmed by quantitative real time PCR there was no overall change in the total Ctnnd1 mRNA levels (Figure 6D), similar to the lack of change in the mRNA levels observed in the DKO mouse model. To further explore, a functional consequence of ESRP1 on protein expression we ectopically expressed Cytoplasmic Esrp1 (ESRP1 -CKLP) and Nuclear Esrp1 (ESRP1 +CKLP) in HEK293T cells transiently transfected with a luciferase reporter bearing the 3’ UTR of the *Gapdh* or the 3’UTR of *Ctnnd1*. We observed that, ESRP1-CLKP caused a modest reduction in the total luciferase activity (Figure 6E), suggesting that ESRP1 functions to inhibit protein translation when bound in the 3’ UTR. This demonstrates a two-tier regulation of CTNND1 by ESRP1, in which ESRP1 regulated splicing and ESRP1 regulated translation control affect the production and expression level of CTNND1 protein isoforms.

## Discussion

Comprehensive analysis of 150 RBPs using eCLIP showed that analysis of RBP binding modalities can infer RBP function (Van Nostrand et al., 2020a; Van Nostrand et al., 2020b). It was shown that RBP binding patterns and cellular localization correlated with function (Van Nostrand et al., 2020b). Many RBPs have multifunctional roles in regulation of RNA processing, and this multifunctionality can result from differences the cellular localization of the RBP (Hafner et al., 2010; Lee et al., 2016; Masuda et al., 2012). We have previously shown that Esrp1 regulates splicing of a large set of epithelial genes and other studies suggest that ESRP1 may also regulate alternative polyadenylation (Bebee et al., 2015; Dittmar et al., 2012).

We generated high resolution RNA map of ESRP1binding sites in the epidermis by integrating peaks from crosslinking immunoprecipitation of ESRP1 from mouse epidermis and analysis of differential alternative splicing between control and *Esrp1/2* deleted epidermis. We confirmed that many ESRP regulated events are directly regulated by ESRP1. Additionally, we provided global evidence supporting the RNA map model wherein ESRP1 binding downstream of the regulated exon promotes exon inclusion, while binding of ESRP1 within or upstream of the regulated exon suppresses exon inclusion. Over 70% of the regulated skipped exon events between control and cDKO epidermis with |ΔPSI| ≥ 20% (quantified by MAJIQ and rMATs using the most stringent criteria) had a peak of ESRP1 near the regulated exon consistent with the position dependent splicing RNA MAP for ESRP1 (Figure 3A, Table S4). While many of the alternatively spliced transcripts directly regulated by ESRP1 had peaks proximal to the regulated exon we observe instances of more distal regulation by ESRP1. We observed several instances of genes with intronic ESRP1 binding peaks that are far (>1kb) to the regulated exon, such as in *Enah, Myo6, Myo9a, and Uap1* (Figure 4A, S3B, S4). In these genes, the peaks of ESRP1 are detected with crosslink sites over 1kb downstream of the regulated exon. Nonetheless, congruent with the position-dependent regulation of alternative splicing of ESRP1 the regulated exons in these genes are upregulated by the presence of ESRP1. In the example of *Enah* exon 11a, the function of a distal splicing enhancer was proposed to involve formation of “RNA bridges” or secondary structures. This distal enhancer contained binding sites for RBFOX2 that, similar to ESRP1, promotes exon splicing. The formation of stem loop secondary structures between the exon and these enhancers was proposed to bring the exons within closer spatial proximity (Lovci et al., 2013). The identification of ESRP1 binding sites within the same region of this enhancer indicates that such stem loops similarly facilitate exon splicing ESRP1 and possibly cooperatively with RBFOX2. Other instances of stem loops within introns bridging genomic distances and facilitating regulated alternative splicing has been shown for additional shared ESRP and RBFOX target genes, such as FGFR2 (Baraniak et al., 2006; Muh et al., 2002). Our findings add further support for the proposal that there are likely thousands of deep intronic splicing regulatory elements (Conboy, 2021). Additionally, we identified ESRP1 binding sites located near tandem alternatively spliced exons, such as those of *Cd44* (Figure S3A), *Myo1b* (Figure 3B), and *Ctnnd1* (Figure S7) providing a mechanism to account for coordinated regulation of these exons.

We observe many large changes in total gene expression upon Esrp ablation, but there is little overlap between these genes and those bound by ESRP1. As the loss of *Esrp1/2* causes an epidermal barrier defect, many if not most of these gene expression alterations, including those of the epidermal differentiation complex, are highly similar to those observed in other mouse models associated with epidermal barrier defects (Bebee et al., 2015; Koch et al., 2000; Segre et al., 1999).. As such we posit that these gene expression changes are secondary to Esrp ablation these genes were not directly bound by ESRP1. Using CLIP we identity robust binding of ESRP1 within the untranslated regions of genes involved in epithelial cell function. We observe peaks for ESRP1 in the CDS and 3’UTR of *Cdh1* and in 5’ UTR, 3’ UTR and CDS of *Ctnnd1* (Figure 6). We previously showed that ESRP1 is associated with CL/P, skin loss and observe hair follicle abnormalities (Bebee et al., 2015; Lee et al., 2018). Further, mutations in CTNND1 and CDH1 have been implicated in the CL/ P (Cox et al., 2018). In the case of CTNND1, we observe that ESRP1 ablation leads to loss of the epithelial isoform of CTNND1, but also an increase in the protein expression of the predominantly mesenchymal isoform of CTNND1 (p120-1) (Figure 6D). It has been shown that while p120-3 is almost exclusively epithelial, the p120-1 isoform, although predominantly mesenchymal, is also expressed in the epithelial cells with lower differentiation or rapid turnover (Venhuizen et al., 2019). The increased protein expression of p120-1 may be attributable to lack of repression by Cytoplasmic ESRP1. One possibility for regulation of CTNND1 by ESRP1, is not only to generate the CTNND1 splice isoform that stabilizes membrane association of CDH1, but ESRP1 may also maintain a balance of these isoforms by repressing the expression of mesenchymal isoforms that associate with other cadherins or that have different functional properties. In epithelial cells CTNND1 is co-localized with CDH1 at adherences junctions and has been proposed to promote cell adhesion and maintain epithelial barriers (Smalley-Freed et al., 2010). This association appears to occur in a tissue specific and isoform specific manner (Venhuizen et al., 2019). In contrast, the mesenchymal CTNND1 isoform has been shown to have opposing functions to promote cell motility and tissue invasion (Yanagisawa et al., 2008). Some of these differences in function of epithelial and mesenchymal CTNND1 isoforms are likely due to differences in protein-proteins interactions as shown for CTNND1 interacting proteins (Markham et al., 2014; Yu et al., 2016) CDH1 is a transmembrane protein and its membrane association is stabilized by CTNND1, however, we have previously shown that loss of ESRPs (which results in a loss of the epithelial CTNND1 (p120-3) isoform) results in diffuse localization of CDH1 in the cytoplasm (Davis et al., 2003; Warzecha et al., 2010).

Overall, we have demonstrated that ESRP1 directly binds and regulates an epithelial splicing network, further that ESRP1 has apparent roles outside of splicing as indicated by its binding within the UTR. While we currently have some evidence in support for a role of ESRP1 in modulating translation, further studies are needed to characterize global changes in protein levels that result from loss of ESRP1. Such studies would include proteomic studies as well as analysis of changes in ribosome association. Another intriguing possibility is that ESRP1 controls mRNA localization of mRNAs with 5’ or 3’ binding sites and localized translation. Previous studies have suggested that mRNA localization in epithelial cells, and localized translation are important for epithelial cell homeostasis (Moor et al., 2017). Given the well-established roles or RNA binding protein binding in 3’ UTRs to regulate RNA stability, it would stand to reason that ESRP1, which is more specifically expressed in epithelial cells than any other RBP, would be uniquely poised to regulate mRNA localization to support epithelial cell functions. Future studies will explore this hypothesis.

## Acknowledgement

We thank Kristen Lynch for critical review of the manuscript, and Fange Liu and Kathryn Hamilton for constructive feedback during manuscript revision. We thank Albert Reynolds for providing us with the p120-Catenin antibodies. We are grateful to the Transgenic and Chimeric Mouse Core of the University of Pennsylvania for generation of Esrp1FLAG/FLAG mice (supported by NIH center grants P30DK050306, P30DK019525, and P30CA016520). We are grateful to the Center for Molecular Studies in Digestive and Liver Diseases (NIH-P30-DK050306) and its core facilities (Molecular Pathology and Imaging Core, Host-Microbial Analytic and Repository Core, Genetically-Modified Mouse Core, and the Cell Culture and iPS Core) for use of luminometer for luciferase experiments. Funding for this work was supported by NIH/NIAMS, 1R56AR066741 (R.P.C), NIH/NIAMS, R01 AR066741 (R.P.C), NIGMS/NIH, P20GM103436 (J.W.P), and NIEHS/NIH, P30ES030283 (J.W.P), and a subproject of a Penn Skin Biology and Diseases Core grant, NIAMS/NIH, 5-P30-AR-057217

## Author Contributions

Conceptualization, N.P and R.P.C., Software, J.Y.H., M.Q.V., P.S., Y.B., and J.W.P., Formal Analysis, N.P., R.P.C., P.S., J.Y.H., and J.W.P., Investigation, N.P., Y.Y., and M.J.S., Resources, Y.B., J.W.P., and R.P.C., Writing – Original Draft, N.P., and R.P.C., Supervision, R.P.C., Funding Acquisition, R.P.C., Writing – Review & Editing, N.P., J.Y.H., M.Q.V., M.J.S., Y.Y., P.S., Y.B., J.W.P., and R.P.C.

## EXPERIMENTAL MODEL AND SUBJECT DETAILS

### Mouse Strains

Generation of Esrp1 KO (*Esrp1*^*-/-*^) and conditional floxed Esrp1 (*Esrp1*^*flox/flox*^) were described previously (Bebee et al., 2015) as was the *Esrp1* ^FLAG/FLAG^ (Lee et al., 2020), and *Esrp1*^*flox/flox*^; *Esrp2*^*-/-*^; *K5rTA tetO-Cre* mice (Lee et al., 2018). Transgenic *Keratin-5 rtTA* (*K5-rtTA*) (Mucenski et al., 2003) mice were obtained from Sarah Millar (Icahn School of Medicine) and *tetO-Cre* strains (Diamond et al., 2000) were obtained from JAX. *Esrp1* deletion in epidermis was induced by Doxycycline (600mg/kg) (Bio-Serv) feeding of nursing mothers at E17.5 until PND5. Genomic DNA for genotyping was derived from tail biopsies and genotyping was performed using standard procedures. Both male and female mice and were used in this study. All animal procedures and experiments were approved by the Institutional Animal Care and Use Committee (IACUC) at the University of Pennsylvania.

### Cell Culture

Py2T (Waldmeier et al., 2012) and HEK293T cells were grown in DMEM with 10% FBS, and Human non-small cell lung cancer cell line H358 (obtained from the American Type Culture Collection) were maintained in RPMI1640 with 10% FBS at 37°C and 5% CO2. *Esrp1* KO Py2T cells were previously described (Lee et al., 2018). H358 knockdown cells are from our previous work (Yang et al., 2016).

## METHOD DETAILS

### Transfections

For luciferase reporter assays, HEK293T cells were transfected (using Mirus Transit 293 reagent) in a 96 well plate with indicated plasmids (total of 20ng DNA) in biological triplicates. Cells were harvested 48hrs after transfections with 100uL Passive Lysis Buffer. Dual Luciferase assay were performed using the Dual-Luciferase Reporter Assay (Promega) according to manufactures instructions. Reporter renilla luciferase activity was normalized to firefly luciferase activity.

### Real-time RT-PCR and RT-PCR

Real-time RT-PCR and RT-PCR were performed as described (Bebee et al., 2015; Lee et al., 2018; Lee et al., 2020). RT-PCR was quantified using ImageLab, Version 6.1 Standard Edition (Biorad). Splicing ratios are represented as PSI for cassette exons and were normalized to RT-PCR product size. Real-time RT-PCR and RT-PCR primer sequences are available.

### Western Blot

Total cell lysates were harvested in RIPA buffer and immunoblotting was performed as described previously (Bebee et al., 2015).Briefly, Total proteins were separated by 4-12% SDS-PAGE and then transferred to nitrocellulose membranes. Membranes were blocked in 5% Non-fat dry milk powder in phosphate buffered saline-tween 20 (PBST), then incubated overnight at 4°C with the primary antibodies. Subsequently, membranes were washed in PBST and incubated for 1 hour with the appropriate secondary antibodies. After washing three times for 10 minutes with PBST, proteins were visualized by chemiluminescent detection (Thermo Scientific).

### RNA extraction

RNA was extracted from the epidermis as described previously (Bebee et al., 2015) with the following modifications. The pups aged P5 were cryoeuthanized and decapitated. Trunk skin was removed and floated dermis side down on 0.25% trypsin/HBSS at 4°C for 16–18 hr. Epidermis and dermis were manually separated using forceps, rinsed HBSS and snap frozen on liquid nitrogen. Epidermis was lysed for RNA isolation in TriZol (Invitrogen, Carlsbad, CA) and RNA was isolated with RNeasy mini kit (Qiagen).

### RNA sequencing and data analysis

Total RNA from epidermis was used for RNA-Seq and samples were processed at Genewiz. Sequencing averaged 147 × 10^6^ reads (range: 104 – 179⨯ 10^6^ reads) per sample. To assess the RNA sequencing data using MAJIQ, adapters were trimmed from RNA-Seq samples using BBDuk, aligned to the mouse GRCm38 genome assembly using STAR v.2.5.1B (Dobin et al., 2013)and sorted and indexed using samtools v.1.9 (Li et al., 2009). Samples averaged 88.93% uniquely mapped reads (range: 87.28 – 90.06%) for the epidermis. For gene expression quantification, salmon v.0.14.0 (Patro et al., 2017) was used in mapping-based mode with selective alignment on trimmed fastq files using GENCODE vM23 annotation to create the index. Differential gene expression analysis was performed with DESeq2 v.1.22.2 (Love et al., 2014). We identified differentially expressed genes by retaining genes that had an adjusted p-value lower than 0.05 and 2-fold difference between genotypes. Differential splicing analysis was performed with MAJIQ v.2.1 using GENCODE vM23 reference transcriptome annotation (Vaquero-Garcia et al., 2016). We identified differentially spliced junctions by retaining junctions that had a delta PSI of at least 10% with a probability of at least 95%. Gene enrichment analysis was performed with EnrichR v.1.0 using a 2018 release of the GO Consortium annotations.

To assess the RNA sequencing data with rMATs, RNA-seq reads were also mapped to the GRCm38.p6 mouse reference genome and the GRCm38.93 transcriptome (Ensembl GTF release 91) using STAR aligner (v2.6.0c) (Shen et al., 2012; Shen et al., 2014). Differential gene expression between the two cell types were calculated using cuffdiff (v2.2.1) (Trapnell et al., 2010), based on the average FPKM at FDR < 5%, > 2-fold when maximum average FPKM from at least one of cell types is larger than 1. Then, to identify differential AS events between the control and conditional knock-out cells, we used rMATS (v3.2.5) that examines all five basic types of AS events (SE, MXE, A5SS, A3SS, and RI). rMATS uses the reads mapped to the splice junctions and the reads mapped to the exon body to estimate the exon usage (PSI: percent-spliced-in (ψ)). The conditional knock-out was compared to the control to identify differentially spliced events with an associated change in PSI (ΔPSI or Δψ) of these events. To compute p-values and FDRs of splicing events with |Δψ| > 0.01% cutoff, we ran rMATS using -c 0.0001 parameter.

### eCLIP and data analysis

#### Library Production

Pups aged P0-P1 were cryoeuthanized and decapitated. Epidermis was isolated as described earlier and after rinsing in HBSS was placed flat on petri dish with HBSS and crosslinked thrice at 400mJ/cm2. The epidermis was minced with a scalpel, and then flash frozen in liquid Nitrogen and stored at -80C until use. Each sample for eCLIP was comprised of the epidermis from two pups (Esrp1^FLAG/FLAG^ or Esrp1^WT/WT^) which were mixed and dounced with a loose pestle in eCLIP lysis buffer. All other processes after lysis were performed as previously described(Van Nostrand et al., 2016).

#### eCLIP sequencing analysis and ESRP1 binding site peak calling using Piranha

eCLIP sequencing for ESRP1 protein by enhanced crosslinking and immunoprecipitation was performed for 6 replicates each for both CLIP and Input. eCLIP sequencing reads from 6 replicates each for both sample group were pooled together, respectively, then trimmed to remove adaptor sequences using Cutadapt (v1.16). The trimmed reads were mapped to the grcm38_snp_tran mouse reference genome using Hisat2 aligner (v2.1.0). For Esrp1 peak calling, we used Piranha (v.1.2.1), a peak-caller for CLIP-seq data. We input the mapped reads from CLIP sample in BED format to Piranha for identifying regions of statistically significant read enrichment having the mapped reads from Input sample as covariate for running Piranha.

#### eCLIP and motif enrichment analysis of ESRP1 binding using PureCLIP and HOMER

Processed read counts from 6 replicate for both CLIP and Input were used to call peaks and crosslink sites using PureCLIP. As PureCLIP does not support analysis on more than 2 replicates, the 6 biological replicates were pooled into 3 replicate pairs and PureCLIP analysis was performed as previously described (Krakau et al., 2017). The HOMER software package was used to identify the motifs four to seven nucleotides in length, enriched on the transcript strand within 10 nucleotides of the cross-link sites and calculate the cross-link densities relative to the identified consensus motif (Heinz et al., 2010).

### Quantification and Statistical Analysis

Statistical analyses of the data were performed in GraphPad Prism. Error bars on qPCR quantification represent upper and lower limits, otherwise, error bars where present represent ± SEM of experiments performed on biological replicates; n ≥ 3 biological replicates. Statistical significance for the Pearson r correlation was calculated using two tailed t-test. Statistical significance where indicated was calculated by Student’s T test and denoted as follows: * = p < 0.01, p < 0.005; **, p < 0.001; ***, p < 0.0001; ****.

### Contact for Reagent and Resource Sharing

Further information and requests for resources and reagents should be directed to and will be fulfilled by the Lead Contact, Russ P. Carstens. Where indicated the requests will be fulfilled with simple Material Transfer Agreement (MTA).

## Supplementary Information

Supplementary Table 1 – MATs_CON-cDKO

Supplementary Table 2 – MAJIQ Heatmap dPSI10 cDKO-Con

Supplementary Table 3 – Piranha Peaks_Splicing

Supplementary Table 4 – Overlapping rMATS and MAJIQ Skipped Exons and CLIP

Supplementary Table 5 – GO Analysis of 3 UTR Binding

Supplementary Table 6 – Resources List – Primers and Plasmids

## Supplementary Figure Legends

Supplementary Figure 1. CLIP Optimizations for the Epidermis (Left) Autoradiogram of crosslinked and non-crosslinked FLAG ESRP1 IP showing RNA: Protein products following limited digestion with RNase I at 0, 5, 20 and 40U. (Right) Western Blot of lysates of FLAG ESRP1 and non-FLAG tagged ESRP1 showing input and immunoprecipitated proteins. Hatched box indicates portion of the membrane extracted for ESRP1:RNA complexes.

Supplementary Figure 2. Analysis of ESRP1 regulation of alternative splicing by MAJIQ and rMATs. A. Number and type of alternatively spliced events identified by MAJIQ between control and experimental epidermis. Putative events only have one junction identified, and could not be attributed to any of the other classes B. Correlation plot of ΔPSI estimated by MAJIQ of 23 skipped exon events validated by RT-PCR, shown in on graph are Pearson Correlation r and p-values. Green dots represent splicing events suppressed by ESRP1 and red dots represent splicing events enhanced by ESRP1. C. Correlation plot of for ΔPSI skipped exon estimated by rMATs and ΔPSI binary skipped exon events estimated by MAJIQ. E. Voila plots showing examples of the detection of alternative last exon events MAJIQ for Gpatch2. Hatched lines indicate splicing event reported in the database but not detected in the RNA-seq, solid lines indicate splicing event detected in the RNA-seq. Red lines show splicing to the ALE exon 9, blue lines show splicing to ALE exon 10 and green lines indicate splicing to ALE exon 16.

Supplementary Figure 3. Complex regulation of alternative splicing by ESRP1. Genome browser views showing ESRP1 binding (CLIP Seq) and transcript levels (RNA-seq) of for *Cd44*(A), *Myo6* (B), and *Fgfr2* (C). A. Alternative splicing of *Cd44* is regulated by ESRP1. Voila plots show differential exon inclusion between Control (CON) and Experimental (cDKO) conditions for *Cd44*, showing ESRP1 enhances inclusion of isoform variable exons 1-10. CLIP-seq densities show CLIP and size matched input, and transcript levels are from RNA sequence in epidermis ablated of Esrps and control epidermis. Individual Crosslink sites identified by PureCLIP are shown for three replicate pairs at top of the browser views, Peaks called by piranha are indicated above the browser view in red. Conservation scores showing Placental mammal conservation by PhyloP at the base. B. Multiple alignment of ∼60 nucleotide region under ESRP1 peak for several vertebrate species shown. Highlighted in green are UGG motifs recognized by ESRP1. Ethidium bromide-stained agarose gel of RT-PCR products of ESRP1 regulated splicing event for *Myo6*. C. Voila plots show differential exon inclusion between Control (Con) and Experimental (Exp) conditions for *Fgfr2*.

Supplementary Figure 4. ESRP1 Enhanced Cassette Exon Events. Browser views of ESRP1 enhanced splicing events for *Atp6v1c2, Ralgps, Slc37a2, Myo9a* and *Uap1* showing ESRP1 peaks located downstream of the regulated exon where it enhances inclusion of the exon. Genome browser views showing ESRP1 binding (CLIP Seq) and transcript levels (RNA-seq). CLIP-seq densities show CLIP and size matched input, and transcript levels are from RNA sequence in epidermis ablated of Esrps and control epidermis. Crosslink sites identified by PureCLIP are shown for three replicate pairs and peaks identified by Piranha are shown at top of the browser views.

Supplementary Figure 5. ESRP1 Silenced Cassette Exon Events. Browser views of ESRP1 enhanced splicing events for *Magi1, Flnb, Scrib*, and *Timm17b* showing ESRP1 peaks located in the upstream intron and/or within the regulated exon where it suppresses inclusion of the exon. Genome browser views showing ESRP1 binding (CLIP-Seq) and transcript levels (RNA-seq). CLIP-seq densities show CLIP and size matched input, and transcript levels are from RNA sequence in epidermis ablated of Esrps and control epidermis. Crosslink sites identified by PureCLIP are shown for three replicate pairs and peaks identified by Piranha are shown at top of the browser views.

Supplementary Figure 6. Splicing RNA Maps for ESRP1. Splicing RNA Map of distribution of ESRP1 CLIP peaks on alternative 3’ splice site (A3SS) (top) and on mutually exclusive exons (MXE) (bottom). Solid blue line shows the ESRP1 CLIP densities over ESRP1 downregulated exons or A3SS, and dotted blue line shows the significance of the peak at each position. Solid red line shows the ESRP1 CLIP densities of ESRP1 upregulated exons or A3SS, with dotted red line indicating significance (p-value) of the binding at the position shown.

Supplementary Figure 7. Regulation of Tandem Exon events by ESRP1. Alternative splicing of *Ctnnd1* is regulated by ESRP1. Voila plots show differential exon inclusion between Control (CON) and Experimental (cDKO) conditions for *Ctnnd1*, showing ESRP1 suppresses inclusion of isoform 1 exons (exons 5 and 6, PSI (Ψ) values representing differential exon inclusion correspond to blue, green and orange lines on the Voila image and the blue, green and orange boxes in the violin plot. Genome browser views showing ESRP1 binding (CLIP Seq) and transcript levels (RNA-seq). CLIP-seq densities show CLIP and size matched input, and transcript levels are from RNA sequence in epidermis ablated of Esrps and control epidermis. Crosslink sites identified by PureCLIP are shown for three replicate pairs at top of the browser views. Conservation scores showing Placental mammal conservation by PhyloP at the base. Ethidium bromide-stained agarose gel of RT-PCR products of ESRP1 regulated splicing event for *Ctnnd1* in control (CON) compared to conditional double knockout epidermis (cDKO or EXP), and in Py2T epithelial cell lines.

