## Supplementary Figures and Legends for "The Global Protein-RNA Interaction map of Epithelial Splicing Regulatory Protein 1 defines a post-transcriptional program that is essential for epithelial cell function"

### Supplement Figure 1

A.

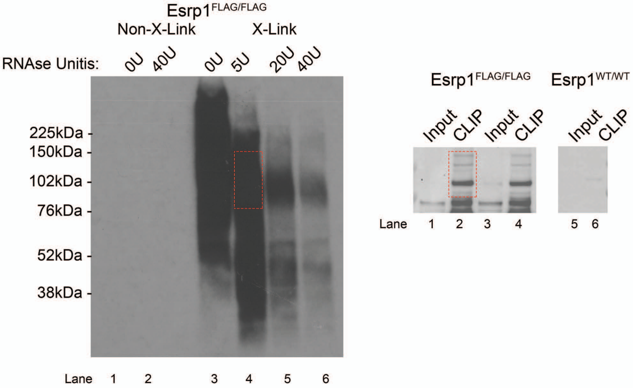

Supplementary Figure 2

A.

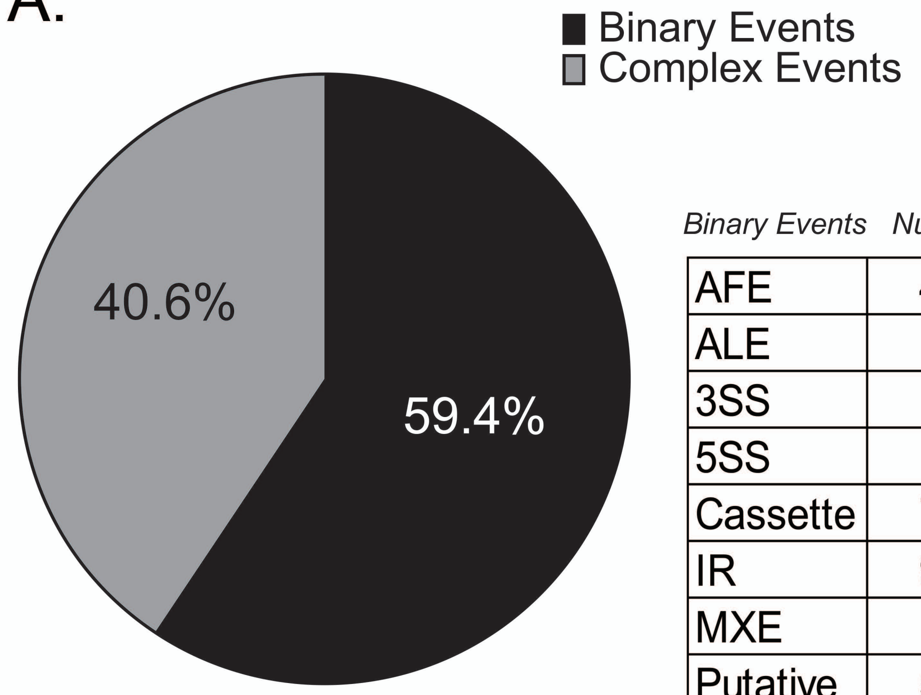


| *Binary Events* | *Number* | *%* |
|---|---|---|
| AFE | 49 | 16.2% |
| ALE | 14 | 4.6% |
| 3SS | 8 | 2.6% |
| 5SS | 8 | 2.6% |
| Cassette | 77 | 25.4% |
| IR | 95 | 31.4% |
| MXE | 2 | 0.7% |
| Putative | 50 | 16.5% |

B.

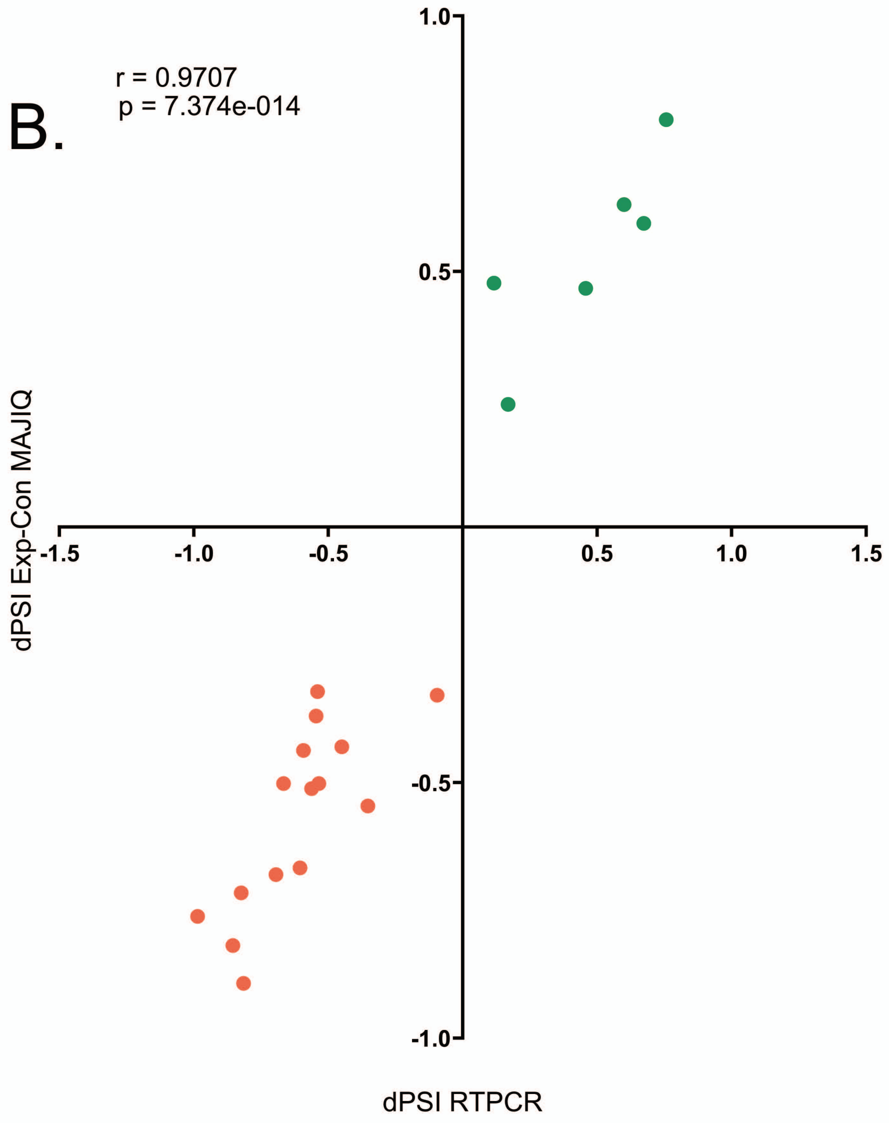


C.

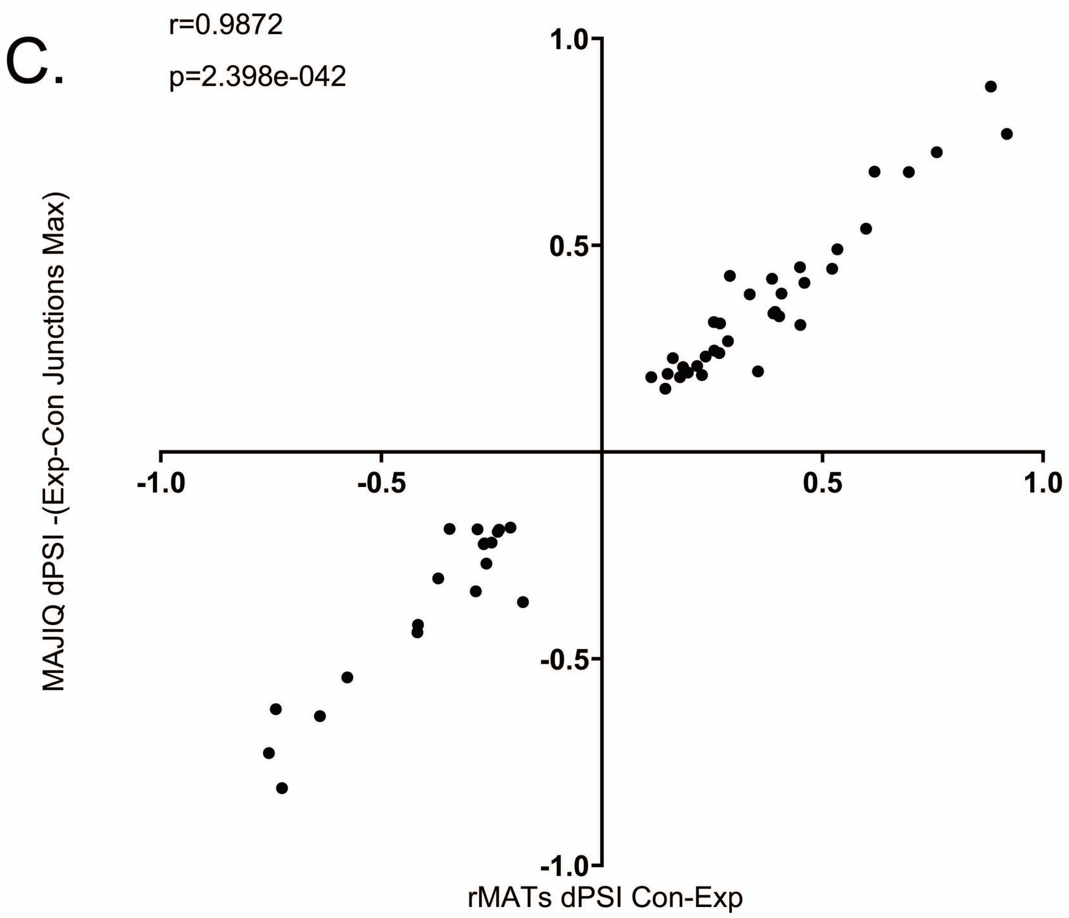


D.

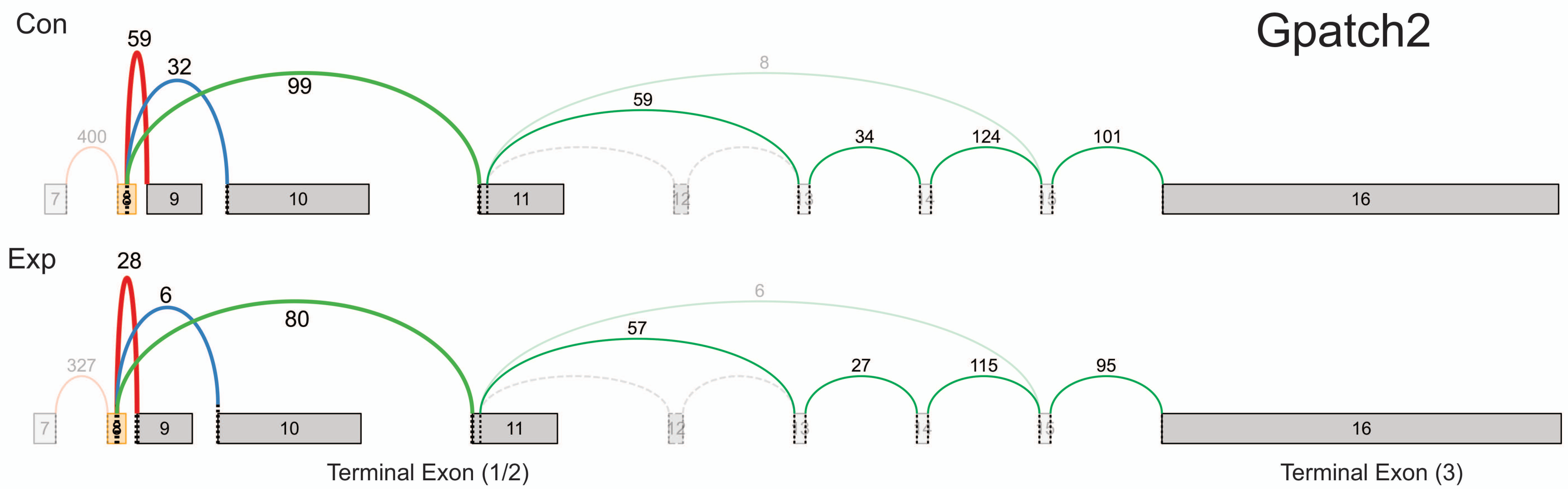

Supplementary Figure 3

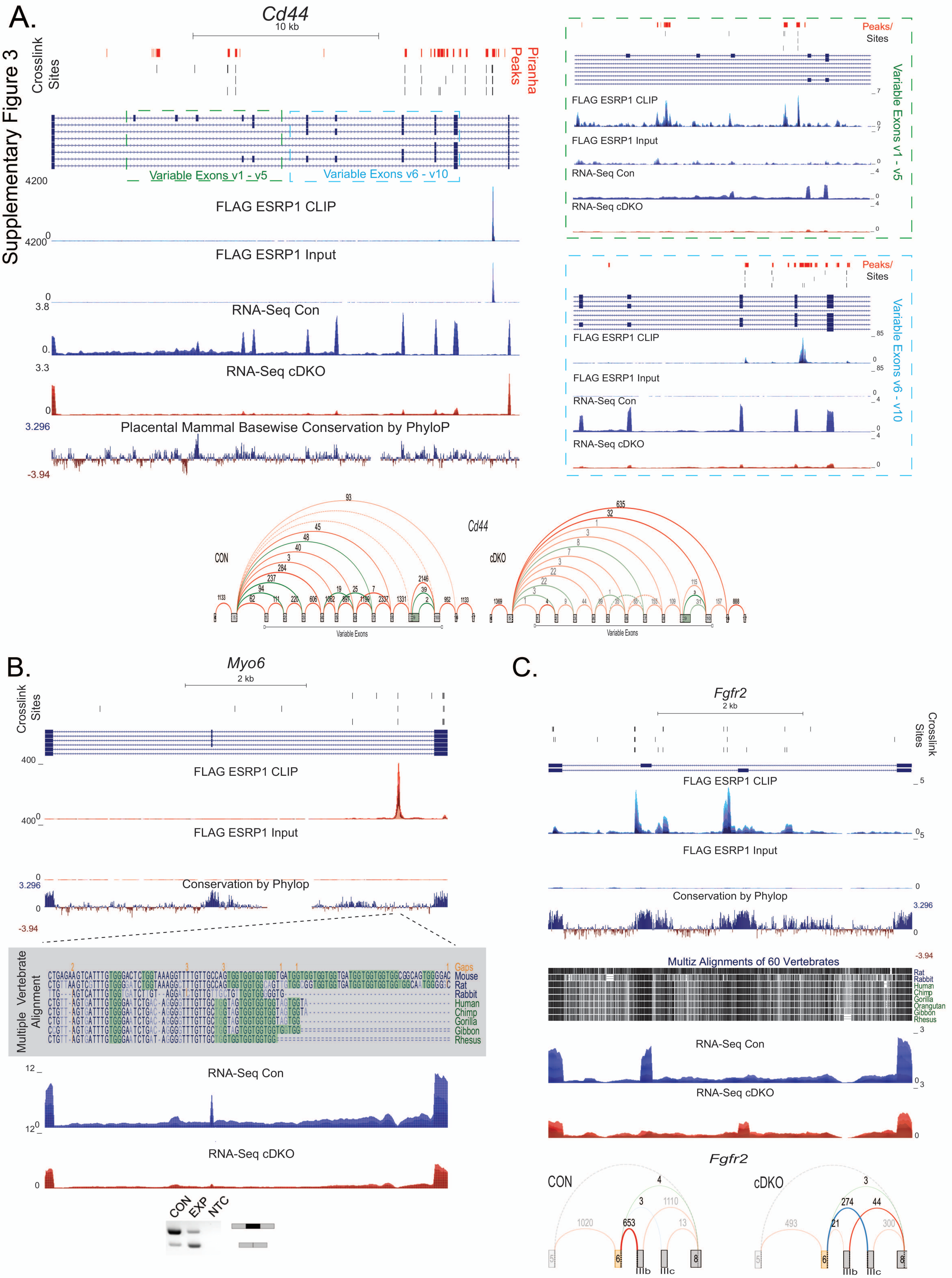

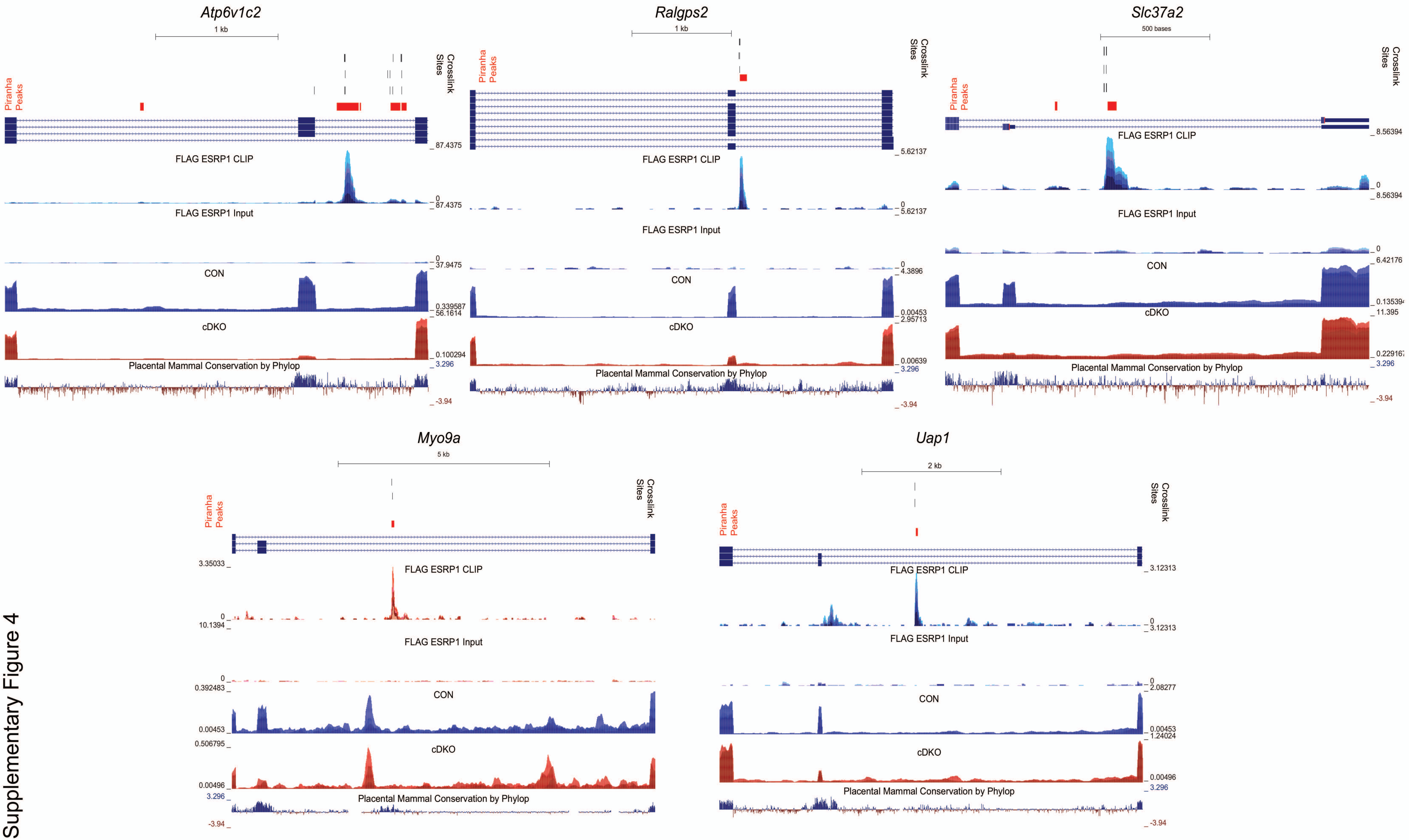


Supplementary Figure 4

Supplementary Figure 5

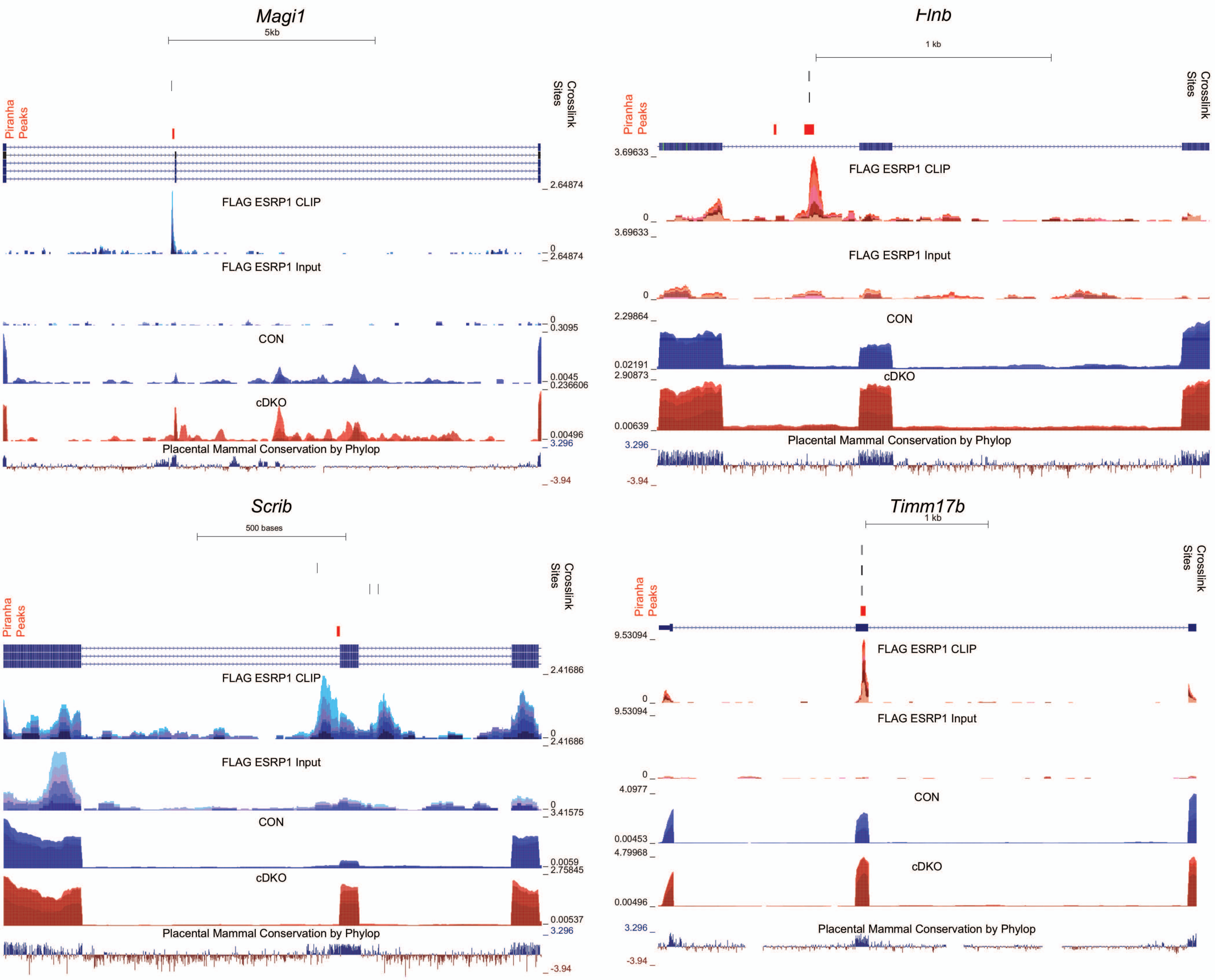

Supplementary Figure 6

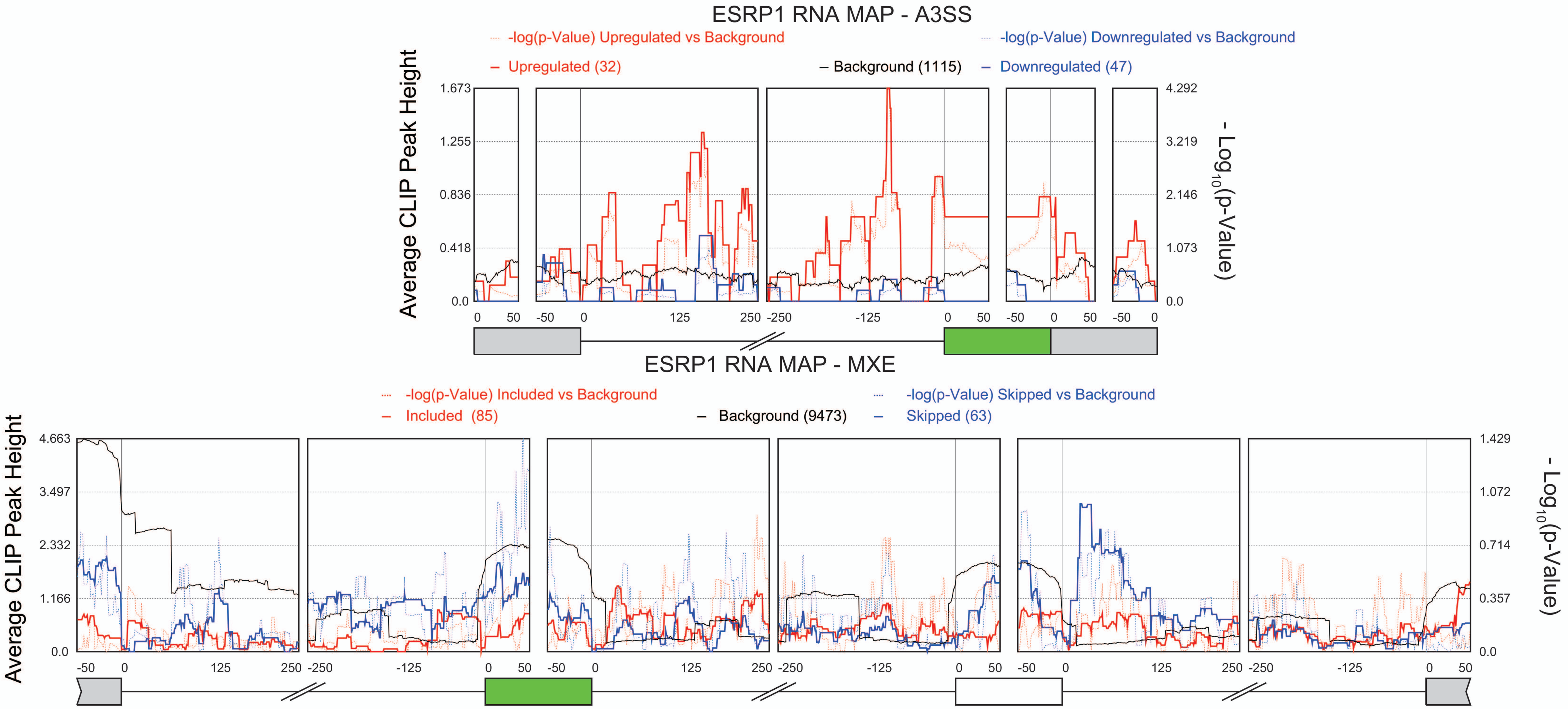

### Supplementary Figure 7

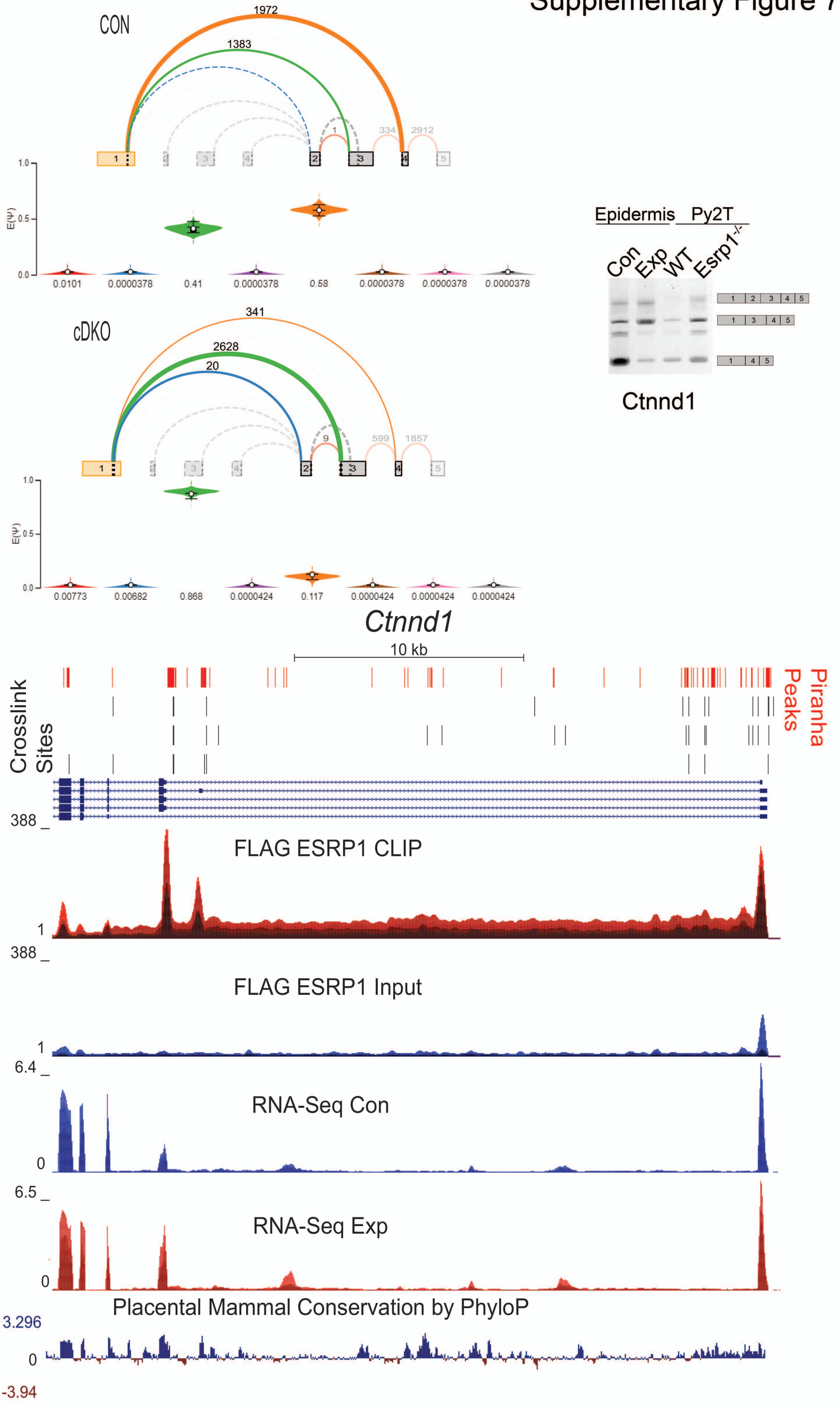

**Supplementary Figure Legends**

Supplementary Figure 1. CLIP Optimizations for the Epidermis (Left) Autoradiogram of crosslinked and non-crosslinked FLAG ESRP1 IP showing RNA: Protein products following limited digestion with RNase I at 0, 5, 20 and 40U. (Right) Western Blot of lysates of FLAG ESRP1 and non-FLAG tagged ESRP1 showing input and immunoprecipitated proteins. Hatched box indicates portion of the membrane extracted for ESRP1: RNA complexes.

Supplementary Figure 2. Analysis of ESRP1 regulation of alternative splicing by MAJIQ and rMATs. A. Number and type of alternatively spliced events identified by MAJIQ between control and experimental epidermis. Putative events only have one junction identified and could not be attribute to any of the other classes B. Correlation plot of $\Delta$PSI estimated by MAJIQ of 23 skipped exon events validated by RT-PCR, shown in on graph are Pearson Correlation r and p-values. Green dots represent splicing events suppressed by ESRP1 and red dots represent splicing events enhanced by ESRP1. C. Correlation plot of for $\Delta$PSI skipped exon estimated by rMATs and $\Delta$PSI binary skipped exon events estimated by MAJIQ. $\Delta$PSI for MAJIQ is reported as cDKO-CON, as such the sign on the values of $\Delta$PSI was changed to reflect the direction of $\Delta$PSI for rMATS (CON-cDKO). E. Voila plots showing examples of the detection of alternative last exon events MAJIQ for Gpatch2. Hatched lines indicate splicing event reported in the database but not detected in the RNA-seq, solid lines indicate splicing event detected in the RNA-seq. Red lines show splicing to the ALE exon 9, blue lines show splicing to ALE exon 10 and green lines indicate splicing to ALE exon 16.
